# Cross-population Joint Analysis of eQTLs: Fine Mapping and Functional Annotation

**DOI:** 10.1101/008797

**Authors:** Xiaoquan Wen, Francesca Luca, Roger Pique-Regi

## Abstract

Mapping expression quantitative trait loci (eQTLs) has been shown as a powerful tool to uncover the genetic underpinnings of many complex traits at the molecular level. In this paper, we present an integrative analysis approach that leverages eQTL data collected from multiple population groups. In particular, our approach effectively identifies multiple independent *cis*-eQTL signals that are consistently presented across populations, accounting for heterogeneity in allele frequencies and patterns of linkage disequilibrium. Furthermore, our analysis framework enables integrating high-resolution functional annotations into analysis of eQTLs. We applied our statistical approach to analyze the GEUVADIS data consisting of samples from five population groups. From this analysis, we concluded that i) joint analysis across population groups greatly improves the power of eQTL discovery and the resolution of fine mapping of causal eQTLs; ii) many genes harbor multiple independent eQTLs in their *cis* regions; iii) genetic variants that disrupt transcription factor binding are significantly enriched in eQTLs (*p*-value = 4.93 × 10^−22^).

**Author Summary:** Expression quantitative trait loci (eQTLs) are genetic variants associated with gene expression phenotypes. Mapping eQTLs enables us to study the genetic basis of gene expression variation across individuals. In this study, we introduce a statistical framework for analyzing genotype-expression data collected from multiple population groups. We show that our approach is particularly effective in identifying multiple independent eQTL signals that are consistently presented across populations in the proximity of a gene. In addition, our analysis framework allows effective integration of genomic annotations into eQTL analysis, which is helpful in dissecting the functional basis of eQTLs.

## Introduction

Expression quantitative trait loci, or eQTLs, are genetic variants that are associated with gene expression levels. Mapping eQTLs can help in dissecting the molecular mechanisms by which genetic variants impact organismal phenotypes. Recent studies [1–3] have revealed that there are substantial overlaps between eQTLs and genetic variants identified from genome-wide association studies (GWAS) of disease phenotypes. In addition, eQTL mapping provides a powerful tool for investigating the regulatory machinery in different tissues [4, 5] or cellular environments [6–8].

In this paper, we *jointly* address three outstanding issues in eQTL mapping. First, due to the high experimental cost, most available eQTL data sets typically have limited sample sizes. To improve power of eQTL discovery, it becomes necessary to aggregate evidence across multiple data sets. Second, because a gene is typically regulated by many regulatory elements, it is highly likely that there exist multiple independent eQTLs in its proximity (i.e., *cis* region). In this scenario, a multi-SNP analysis is required to uncover all relevant *cis* acting genetic factors involved in the gene regulation process [9]. Third, the availability of extensive functional annotations [10–12] now enables integration of functional genomic information into eQTL analysis, which can be useful to dissect the functional basis of eQTLsf. Linking genomic annotations to eQTLs goes beyond genetic association analysis, and helps gain a better understanding of the underlying biological processes. Individually, some of these three issues have been discussed by previous works. For example, [3, 9, 13–16] discussed single SNP analysis of eQTLs jointly from different studies, populations or tissues. But these methods do not naturally extend to multi-SNP analysis. [17–20] examined the enrichment of selected genomic features in *cis*-eQTLs, mostly based on single SNP association results. To the best of our knowledge, there is no existing approach that jointly addresses all three issues in a systematic way.

In this paper, we demonstrate an integrative analysis framework to perform multi-SNP fine mapping analysis of eQTLs using cross-population samples. Our statistical methods stem from an established Bayesian framework proposed by [14, 15, 21], which has been successfully applied in mapping eQTLs from multiple tissues. We apply our statistical framework to analyze the data from the GEUVADIS project [20], where the expression-genotype data are collected from five population groups. In GWAS, trans-ethnic meta-analysis of genetic association data from diverse populations has been shown to be a powerful tool in detecting novel complex trait loci and improving resolution of fine mapping of causal variants by leveraging population heterogeneity in local patterns of linkage disequilibrium (LD) and allele frequencies [22, 23]. This approach, to the extent of our knowledge, has not been applied to eQTL analysis. Utilizing cross-population expression-genotype data, we are interested in identifying eQTL signals that show *consistent* effects in all populations. Furthermore, we aim to examine whether we have sufficient statistical power to identify multiple independent *cis*-eQTL signals with the available aggregated sample size. Finally, we set out to investigate whether the genetic variants that disrupt transcription factor (TF) binding are enriched in eQTLs, and we integrate such annotations to further improve fine mapping analysis of eQTLs. Our three main aims are also inherently inter-related and reinforce each other. With higher power through sample aggregation in mapping eQTLs, our method can identify at a high resolution potentially multiple genomic regions that harbor casual eQTLs. Consequently, these efforts improve the statistical power and precision of localization for our functional analysis.

## Results

### Method Overview

We start with a brief description of our statistical framework and general strategy for multi-SNP fine mapping analysis in a meta-analytic setting across multiple populations.

#### Statistical Model and Inference

Consider a genomic region with *p* SNPs that is interrogated in *s* different population groups. In each group *i*, we use a multiple linear regression model to describe the potential genetic associations between the *p* SNPs and the expression levels of a target gene:

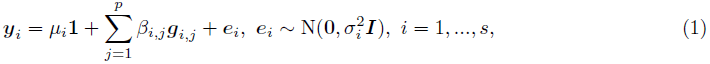

where the vectors *y*_*i*_ and *e*_*i*_ represent the expression levels and the residual errors in population group *i*, and the parameters *μ*_*i*_ and 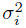 denote the intercept and the residual error variance specific to the population group. The vector ***g***_*i, j*_ denotes the genotype of SNP *j* in population group *i*, and the regression coefficient *β*_*i, j*_ represents its genetic effect. Across all population groups, the *s* linear models form a system of simultaneous linear regressions (SSLR, [15]).

The problem of mapping eQTLs can be framed as identifying SNPs with non-zero *β*_*i, j*_ values. For each SNP *j*, we define an indicator vector

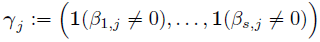

to represent its association status in each of the *s* population groups. Such indicator is referred to as a “configuration” in the literature of genetic association analysis across multiple subgroups [14, 15, 24]. For each target gene, our computational procedure is designed to make joint inference on all *p* SNPs with respect to 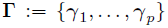 given observed genotype data, 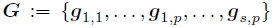, and expression phenotype data, 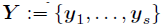.

For mapping eQTLs in a cross-population meta-analytic setting, we make an important assumption that if a SNP is genuinely associated with a given expression phenotype, its underlying genetic effects are non-zero in *all* population groups. That is, *γ*_*j*_ = **1**, if SNP *j* is an eQTL, and **0** otherwise. This assumption is largely motivated by the biological hypothesis that the regulatory mechanisms behind eQTLs likely remain the same across populations. Here, we acknowledge that there exists convincing evidence for population specific eQTLs [20], however, by employing the above assumption, our analysis focuses on identifying eQTLs whose effects are consistent across populations, which are the vast majority of the cases [21].

For each SNP *j*, we assume an independent prior for *γ*_*j*_, which assigns most of the probability mass on *γ*_*j*_ = **0** and encourages an overall sparse structure of **Γ**. This is because most previous studies only identified small numbers of *cis*-eQTLs for any given gene. Particularly in our fine mapping analysis where *p* is typically in the magnitude of 10^3^ to 10^4^, we use

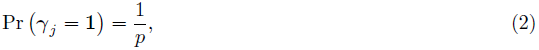

which implies that we expect a single *cis*-eQTL signal for the target gene *a priori*, as

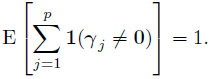

We will further justify this prior specification based on our overall strategy for fine mapping analysis, and discuss some other alternative prior specifications for enrichment analysis in the later sections.

Conditional on SNP *j* being an eQTL, we model its genetic effects across populations, 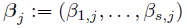,using a flexible Bayesian prior for meta-analysis proposed in [21]. Briefly, we assume that the effect sizes of an eQTL are highly correlated while allowing some reasonable degree of heterogeneity across population groups (See the Material and Methods section for details).

Given observed data (***Y, G***) and a specified **Γ** value, a Bayes factor

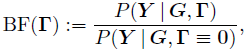

can be analytically approximated with high accuracy following [15]. Based on this result, it is straight-forward to compute the posterior of **Γ** using the Bayes rule, i.e.,

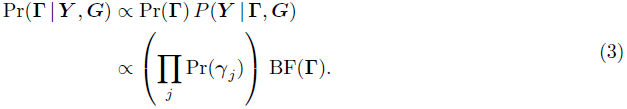

We implement an efficient MCMC algorithm to perform the posterior inference and summarize the fine mapping results from the posterior samples. More specifically, we compute the posterior probability for each possible **Γ** by the corresponding frequency in the posterior samples. We will refer to this quantity as “posterior model probability” henceforth. To evaluate the importance of each SNP, we compute a posterior inclusion probability (PIP) for each SNP by marginalizing over all posterior model probabilities. If a SNP is included in posterior models with high frequencies, its corresponding PIP tends to be large.

#### Dealing with Genetic Data from Multiple Populations

We note that analyzing eQTL data using samples collected from multiple populations is slightly different from the standard settings in meta-analysis of genetic associations. In particular, the allele frequencies of interrogated SNPs and/or patterns of linkage disequilibrium presented in genotype data can be highly heterogeneous in different populations. The proposed multi-SNP fine mapping procedure takes advantage of the unique setting of cross-population samples, and leverages statistical power for eQTL discovery.

The allele frequency of a SNP is not directly connected to its underlying genetic effect with respect to expression levels. However, because of its impact on sampling errors, it affects the precision of *estimated β*_*i, j*_ (denoted by 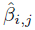) in model (1). Lower allele frequency typically results in larger uncertainty (e.g., the standard error of 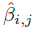 of a rare SNP is usually larger than that of a common SNP), which implies that the estimates of rare SNPs tend to be noisy and less informative. In the extreme case if a SNP is monomorphic in samples from a particular population, it should be considered completely uninformative for examining genetic associations. As shown by [15], Bayesian procedures, especially in use of Bayes factors, can precisely capture the (non-)informativeness of SNPs in each population group: e.g., including the genetic data of monomorphic SNPs yields identical inference results as discarding such SNPs from corresponding population groups. On the other hand, our adopted Bayesian meta-analysis prior enables aggregating less informative or weak association signals from low frequency SNPs, as long as they are *consistent* across population groups.

In pursuing consistent association signals, our fine mapping procedure takes advantage of potential varying LD patterns across multiple population groups. This is mainly because our method favors identifying eQTLs whose effects are consistent in *all* population groups, whereas SNPs that tag causal variants only in *some* populations (due to population-specific LD structures) are automatically down-weighted. As a consequence, the genomic regions that harbor causal eQTLs can be effectively narrowed down using cross-population data. This advantage becomes even more obvious when performing *multiple* SNP analysis, as all candidate *cis*-SNPs are simultaneously evaluated.

#### Gene-level Testing Prior to Fine Mapping *cis*-eQTLs

In most currently available data sets, eQTL discoveries are only confidently made in a proportion of genes. To reduce the computational cost of the fine mapping analysis, we adopt a practical procedure that first screens the genes having at least one *cis*-eQTL (which we will call “eGenes”).

The statistical procedure to identify eGenes across multiple subgroups has been well established in [14]. More specifically in the context of cross-population eQTL mapping, we assume a similar linear model system as (1), and perform gene-level Bayesian hypothesis testing. Namely, for each gene, we test

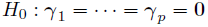

versus

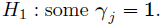

The eGenes are identified upon rejecting the corresponding null hypotheses, and we only follow up the eGenes with the multiple *cis*-eQTL analysis.

The gene-level testing is computationally efficient, and it effectively filters out a substantial set of genes that are unlikely to be interesting for fine mapping. Furthermore, this additional procedure justifies our use of prior (2): given that a gene is identified as an eGene, expecting a single *cis*-eQTL prior to the fine mapping analysis seems, on average, a slightly conservative assumption.

#### Enrichment Analysis and Integration of Genomic Features

We further consider incorporating additional SNP-level functional annotations in our multiple *cis*-eQTL analysis to address the following two closely related problems:

1. assessing the enrichment of certain functional annotations in eQTL signals
2. integrating the enriched functional annotations to improve the resolution of fine mapping analysis of eQTLs

The first problem of enrichment assessment can be framed as a hypothesis test of correlations between the (eQTL) association status of all *cis*-SNPs (*γ*_*j*_’s) and their functional annotations. Note, to properly quantify the level of enrichment, it is required to jointly analyze all available gene-SNP pairs instead of focusing only on eGenes. The intrinsic statistical challenge lies in the fact that the values of *γ*_*j*_’s are not directly observed and the inference is required from the expression-genotype data. Here, we propose to test the association between the PIPs from the output of the multi-SNP *cis*-eQTL analysis, Pr(*γ*_*j*_ | ***Y, G***), and the genomic annotations of interest in a logistic regression model framework. More specifically, we first carry out the proposed Bayesian multi-SNP analysis to compute PIPs for *all* gene-SNP pairs using the following prior specification,

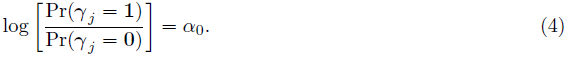

In particular, we determine *α*_0_ in a similarly conservative way as (2) using the results from gene-level testing as follows: Let *g*_*e*_ and *g*_*s*_ denote the number of eGenes identified and the number of total gene-SNP pairs in the analysis, respectively. We set

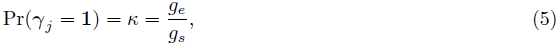

which essentially assumes that each eGene contributes only a single associated gene-SNP pair, and corresponds to 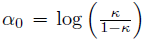. We then fit a logistic regression model by correlating the resulting PIP with the genomic annotations of each SNP and controlling for other potential contributing factors. The key idea is that, under the described settings, the PIPs are computed assuming the null model of no enrichment of any sort. As a consequence, a strong association between the PIPs and the annotations of interest presents evidence against the null hypothesis.

Upon successful enrichment assessments of genomic annotations, our Bayesian framework provides a natural way to integrate relevant annotations into the fine mapping analysis. In particular, we use a generalization of prior specification (4) for Pr(*γ*_*j*_) to incorporate the genomic annotations of SNP *j* with respect to the target gene, i.e.,

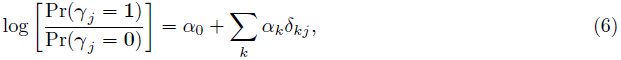

where *δ*_*k j*_ denotes the *k*-th annotation for SNP *j*. The regression coefficient *α*_*k*_, which we refer to as “enrichment parameter”, represents the strength of association between the *k*-th genomic annotation and the prior probability of a SNP being an eQTL. Our inference procedure based on the Bayesian hierarchical model with prior specification (6) involves estimating both the enrichment parameters *α*_*k*_’s and *γ*_*j*_’s in an iterative fashion. We give the details of this procedure and its statistical justification in the Materials and Methods section.

### Simulation Studies

We performed a series of simulation studies to evaluate the proposed Bayesian multi-SNP fine mapping procedures. Specifically, we simulated under the setting where eQTL data are collected from multiple population groups. This section briefly summarizes the main simulation results, and we leave the relevant details in the Materials and Methods section.

First, we examined the power of the proposed Bayesian multi-SNP analysis approach in identifying multiple independent association signals using cross-population samples. In particular, we compared its performance with the commonly applied single SNP meta-analysis procedure and a (step-wise) conditional meta-analysis procedure [25, 26], treating each population group as a participating study in a meta-analysis. To simulate the genotype-expression data in multiple population groups, we used the real genotypes of 2,500 SNPs from 100 randomly selected and distinct genomic regions (i.e., 25 consecutive SNPs per region) in the GEUVADIS data. As a consequence, there are generally modest to high levels of LD within each region, but unremarkable levels of LD between regions. (In other words, we assembled 100 relatively independent LD blocks for each gene.) It is worth noting that within each region, the variation in patterns of LD between populations is maintained. We randomly assigned one to four eQTL SNPs for each gene and simulated their expression levels according to a system of linear models (see Material and Methods section and Section S.5 of S1 Text). For evaluation, we assessed the ability of each procedure in correctly identifying the 25-SNP regions (i.e., the LD blocks) that harbor the true eQTLs. The simulation results indicate that the proposed Bayesian multi-SNP analysis approach is the most powerful in identifying independent eQTL signals among the three methods compared (Fig 1). Further investigation reveals that all three methods achieve similar power when genes contain only a single eQTL; however when a gene harbors multiple independent *cis*-eQTLs, the proposed Bayesian fine mapping approach becomes highly advantageous (Fig 11). In addition, the simulation results show that the reported PIPs from the proposed Bayesian approach are well-calibrated (Fig 13).

**Fig. 1.**
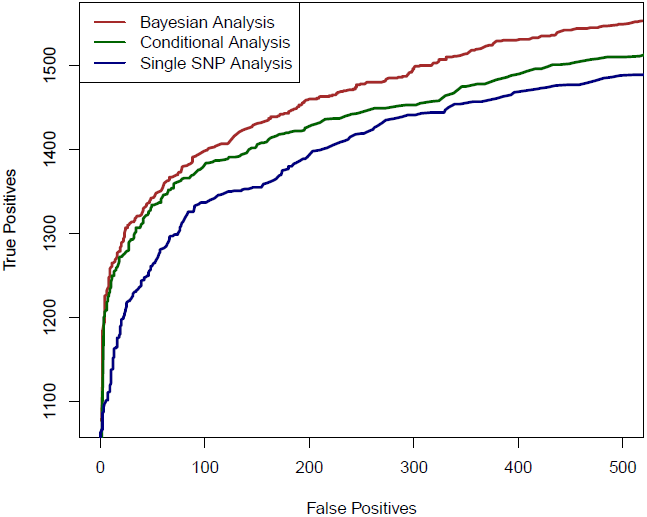
The comparison of powers in identifying independent causal eQTL regions. We compare the performance of the three competing methods in determining the regions harboring the true causal eQTL in a cross-population meta-analytic setting: the proposed Bayesian multi-SNP analysis method (brown line), a conditional meta-analysis approach (dark green line) and a single SNP meta-analysis approach (navy blue line). Each plotted point on the figure represents the number of true positive findings versus the number of false positive findings of a given method at a particular threshold. For any false positive value, the proposed Bayesian approach always yields the most true positive findings.

We also performed additional simulations to evaluate the proposed computational procedures for enrichment testing and fine mapping analysis integrating genomic features. In brief, we found that the proposed procedures have greatly improved power in testing enrichment of genomic features when compared with standard approaches based on single SNP testing results (Table 14). In addition, we observed that the proposed computational procedure provides accurate estimates of the enrichment parameters (Fig 14).

### Analysis of GEUVADIS Data

In this paper, we focused on analyzing the expression and genotype data collected from the GEUVADIS project [20]. More specifically, the data set consists of RNA-seq data on lymphoblastoid cell line (LCL) samples from five populations: the Yoruba (YRI), CEPH (CEU), Toscani (TSI), British (GBR) and Finns (FIN). In our analysis, we selected 420 samples which were densely genotyped in the 1000 Genomes Phase I data release [27] and 11,838 protein coding genes and lincRNAs that are deemed expressed in all five population groups. Throughout, our analysis focused on the SNPs that locate within a 200kb genomic region centered at the transcription start site (TSS) of each gene. In contrast to the original eQTL mapping approach discussed in [20], we treated each population as a single group and performed *cis*-eQTL analysis jointly across all five groups. We have made the full analysis results available on the website http://www-personal.umich.edu/~xwen/geuvadis/.

**Tab. 1.** Comparison of type I error rate and powers between the proposed Bayesian enrichment testing method and a standard enrichment testing procedure

| Method | Type I error | Power |  |  |
| --- | --- | --- | --- | --- |
| | $\alpha_1 = 0$ | $\alpha_1 = 0.25$ | $\alpha_1 = 0.50$ | $\alpha_1 = 0.75$ |
| Multi-SNP PIP | 0.02 | 0.28 | 0.62 | 0.86 |
| *p*-value Classification | 0.01 | 0.12 | 0.17 | 0.33 |

The Bayesian method utilizes PIPs from the multi-SNP eQTL fine mapping analysis. The approach in comparison ranks the SNPs by their single SNP testing *p*-values in each gene, and classifies the top associated SNPs as the “causal” eQTLs if their *p*-values pass the significance threshold at FDR 5% level. Both methods (conservatively) control the type I errors at desired 5% level. The PIP based method show much improved powers comparing to the standard approach in all three alternative scenarios where the magnitude of enrichment ranges from small to modest.

#### Power Gain in gene-level Meta-Analysis

We started our analysis of the GEUVADIS data by performing gene-level testing jointly across all five population groups.

In total, we identified 6,555 eGenes from 11,838 tested protein coding and lincRNA genes at 5% FDR level. For comparison, the numbers of eGenes identified using each population data alone are given in Table 2. The separate analysis identifies no more than 2,100 eGenes in any of the population groups. The union of the eGenes from the separate analysis yields 3,447 genes, the vast majority of which (except for 60 genes) are included in the discoveries by the joint meta-analysis. In addition, the meta-analysis identifies a total of 3,168 new eGenes.

Examining the set of eGenes uniquely identified in the meta-analysis, we found that they share the following common feature: when performing separate analysis within each population group, the strongest association signal in each respective *cis*-region only shows modest strength and does not pass the required significance threshold; however, across population groups, the same association signal tends to be highly consistent, and the overall evidence aggregated across population groups becomes quite strong. As a consequence, the joint analysis of all population groups is able to detect such signals. We demonstrate one of such examples in Fig 2 using gene *NME1* (Ensembl ID: ENSG00000239672), where the gene-level Bayes factor in the joint analysis is several orders of magnitude higher than the corresponding gene-level Bayes factors in each separate population group analysis.

**Fig. 2.**
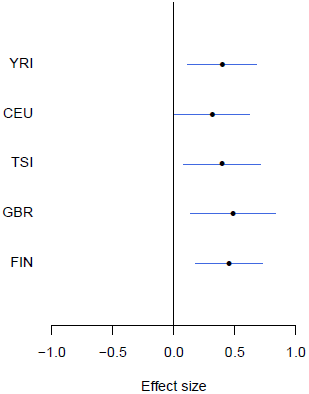
An example of modest yet consistent eQTL signals across population groups. The forest plot shows the genetic effects of SNP rs7207370 with respect to the expression levels of gene *NME1* (Ensembl ID: ENSG00000239672). SNP rs7207370 is one of the top associated *cis*-SNPs in all population groups, yet the strengths of the association signals are modest in all groups (the maximum single SNP Bayes factor, among five groups, is 18.0 in FIN, the corresponding gene-level Bayes factor is 1.8). As a consequence, the gene is not identified as an eGene in any of the separate analyses. Across populations, the SNP exhibits a strongly consistent association pattern. In the cross-population meta-analysis, the gene-level Bayes factor reaches 1.1 × 10^4^ (single SNP Bayes factor for rs7207370 is 9.5 × 10^5^).

**Tab. 2.** Comparison of eGene Discovery by Separate vs. Meta Analysis

| Population | Sample Size | Number of eGenes |
| --- | --- | --- |
| YRI | 77 | 1042 |
| CEU | 78 | 960 |
| TSI | 92 | 1803 |
| GBR | 84 | 2078 |
| FIN | 89 | 2100 |
| META | - | **6555** |

eGenes are declared by rejecting the null hypothesis of no *cis*-eQTLs in the *cis* region at 5% FDR level. The difference of identified eGenes between CEU, YRI and the rest of the European populations in the separate analysis is likely due to the cell line effects (T. Lappalainen, personal communication). Clearly, the meta-analysis improves power of eGene discovery by aggregating samples across population groups. In comparison, the union of the eGenes from the separate analysis contains only 3,447 genes.

#### Multi-SNP analysis of eGenes

We followed up the gene-level analysis by performing multi-SNP fine mapping for the set of identified eGenes across all five population groups.

One of our primary aims is to identify potential multiple independent *cis*-eQTL signals while accounting for varying LD patterns in different populations. First, we asked how often we can identify multiple *cis*-eQTL signals in eGenes. To this end, for each fine mapped eGene, we computed the expected number of independent *cis*-eQTL signals from the corresponding posterior distributions. Fig 3 shows the histogram of posterior expected *cis*-eQTL signals in all 6,555 eGenes. It is clear that for the available (accumulated) sample size in the GEUVADIS data, we identified only single eQTL signals for the majority of the eGenes. Nevertheless, for a non-trivial proportion of genes, there is strong evidence that multiple *cis*-acting regulatory genetic variants co-exist and can be confidently identified. More specifically, there are about 14% of the eGenes (or 7% of all interrogated genes) with the posterior expected number of *cis*-eQTLs ≥ 2.

**Fig. 3.**
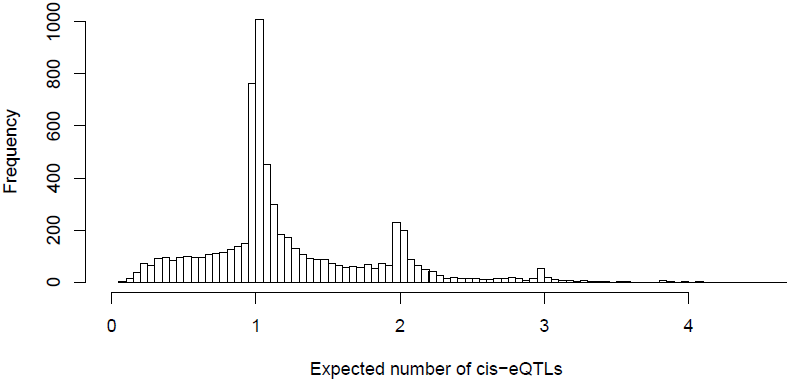
Histogram of posterior expected number of *cis*-eQTLs in 6,555 identified eGenes. The figure indicates that we identified only single *cis*-eQTLs for most eGenes. However, for a non-trivial proportion of eGenes, multiple independent eQTL signals were identified.

For example, in the case of gene *LHPP* (Ensembl ID: ENSG00000107902), four independent eQTL signals are confidently identified (Fig 4). Each eQTL signal (except one) is represented by a cluster of highly correlated SNPs in a small genomic region. Due to LD, within each cluster the correlated SNPs tend to have similar PIPs and we cannot be certain which SNP is truly driving the association signal. However, the sum of the PIPs within each cluster is very close to 1, indicating near certainty that an eQTL is located within the region. There were 134 different posterior models examined for gene *LHPP* in the sampling phase of the MCMC run. Interestingly, every model contains exactly 4 SNPs (which results in posterior expected number of *cis*-eQTLs being 4 with variance 0), each from one of the four independent clusters.

**Fig. 4.**
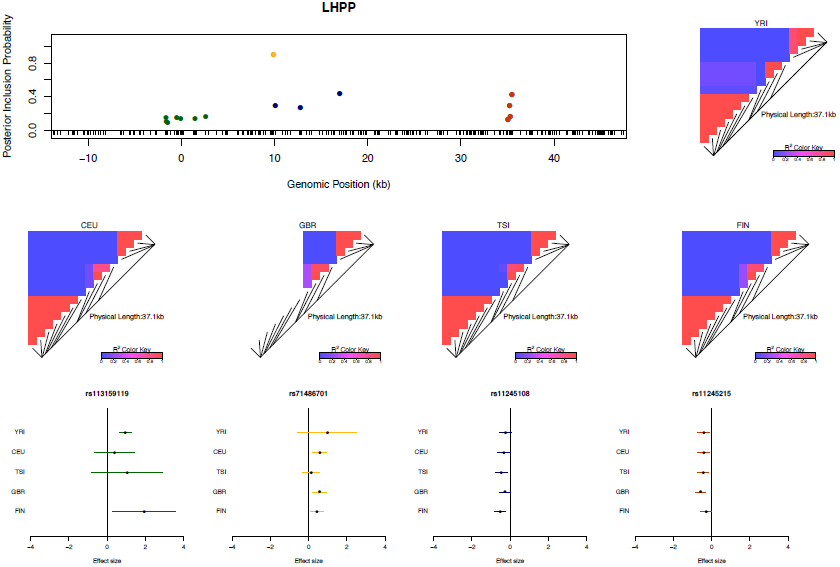
An example of a gene harboring four independent *cis*-eQTL signals. The top left panel plots the *cis*-SNPs with PIP ≥ 0.02. The locations of the SNPs are labeled with respect to the TSS of gene *LHPP*. The ticks on the x-axis indicate all interrogated *cis*-SNPs in the region. The SNPs with the same color are in high LD and represent the same eQTL signal. In the plot, the sums of the PIPs from the SNPs in the same colors are all ~ 1, indicating that we are confident of the existence of each signal. The heatmaps show the LD patterns in each of the population group. They are qualitatively similar, except that the SNPs representing the first signals are monomorphic in GBR. In the bottom panel, we plot the effect sizes of eQTLs jointly estimated from one of the high posterior probability models. Each of the SNP plotted belongs to a different colored cluster in the PIP plot (as indicated by the color coding of the error bars).The effect sizes and standard errors are estimated from the multiple linear regression models (containing all four SNPs) separately fitted in each population group. All the signals show strong effect size consistency across populations.

As previously discussed, even in the cases when only a single *cis*-eQTL signal is identified, we observed that the fine mapping procedure takes advantage of varying LD patterns across populations and narrows down the set of candidate causal variants by down-weighting SNPs that tag causal variants only in some populations. For example, in analyzing the data of gene *AGO3* (Ensembl ID: ENSG00000126070) from TSI alone, we identified a strong single eQTL signal within a 144kb region. The region contains 41 SNPs that are in perfect LD and show equal strength of associations. In the other four populations, this particular region is broken into smaller LD blocks and the associations for the 41 SNPs become distinguishable. As a result, when performing multiple-SNP analysis across all populations, we narrowed down the potential causal eQTL into a 1.2kb region, with only 3 candidate SNPs (in high LD in all populations) fully explaining the observed association (Fig 5).

**Fig. 5.**
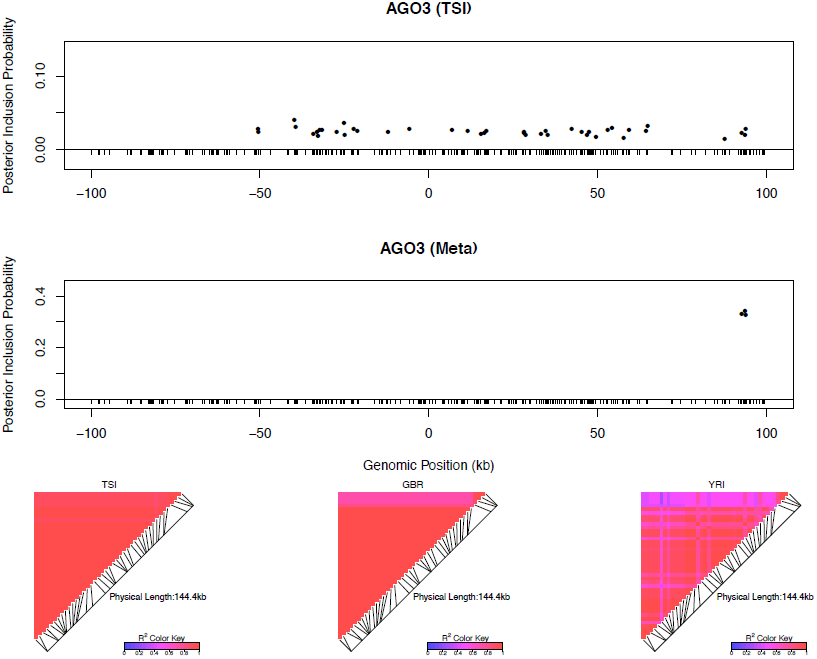
An example of automatic LD filtering across population groups. The top panel shows the result of multiple *cis*-eQTL analysis for gene *AGO3* using only the data from TSI. The SNPs with PIPs ≥ 0.02 are plotted. All SNPs plotted are in high LD in TSI, and the sum of the PIPs across the genomic region is close to 1. The region spanned by the signals is ~ 140 kb. We repeated the analysis jointly across all five populations, the SNPs with PIPs ≥ 0.02 are plotted in the middle panel. The genomic region harboring the eQTL is narrowed down into a 1.2 kb region enclosed by three SNPs each with PIP ~ 0.33. The bottom panel shows the LD heatmaps between the 41 SNPs plotted in the top panel in TSI, GBR and YRI, respectively.The multiple *cis*-eQTL mapping method takes advantage of the varying LD patterns across populations, and automatically narrows down the region harboring the true causal *cis*-eQTL.

To further quantify the effect of LD filtering, we selected a set of 526 eGenes that are highly likely to harbor exactly one *cis*-eQTL based on our multi-SNP analysis. More specifically, we selected the genes whose posterior expected number of *cis*-eQTLs = 1 and variance ≤ 1 × 10^−4^. We then ran multiple *cis*-eQTL analysis separately in each of the five populations for each selected gene. For every multi-SNP analysis of each gene, including the meta-analysis, we constructed a 95% credible set that contains the minimum number of SNPs with the sum of PIPs ≥ 0.95, and defined its credible region length as the distance between the right-most and left-most SNPs in the set. The histogram of reduction in credible region length by cross-population analysis is shown in Fig 6. In 486 out of 526 (or 92%) genes, we found that the cross-population analysis yields the smallest credible region length. The median ratio of credible region length by the joint analysis to the minimum credible region length by the corresponding separate analyses is 0.50 across all 526 genes, indicating that, on average, cross-population meta-analysis can effectively narrow down the genomic regions that harbor *cis*-eQTLs.

**Fig. 6.**
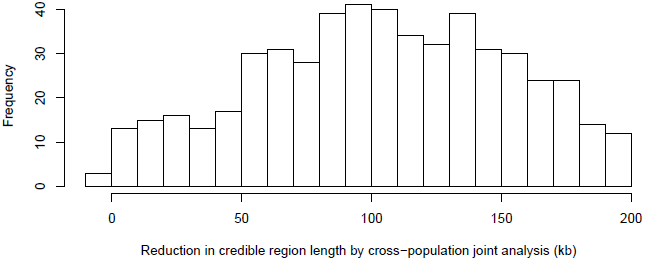
Histogram of reduction in 95% credible region length by cross-population joint analysis. The histogram shows the differences in the credible region lengths between the average of the five separate population analyses and the joint analysis. The analysis is performed using the set of 526 eGenes that highly likely harbor exactly one *cis*-eQTL. Only 3 out of 526 eGenes show (slightly) increased credible region lengths (negative reduction values) in the joint analysis.

Multi-SNP analysis can also be very helpful in explaining some of the extreme heterogeneity of eQTL effects across populations observed in *single* SNP analysis. In particular, we identified a few SNPs that, when analyzed alone, appear to show strong but opposite genetic effects on expression levels in different populations. It seemingly suggests that a particular allele of the variant increases expression levels of the target gene in one population and decreases expression levels in another population. However, going through all such individual examples, we found that none of the opposite effect association patterns is supported by our multi-SNP analysis. In mapping *cis*-eQTLs for gene *TTC38* (Ensembl ID: ENSG00000075234), we found a set of tightly linked SNPs displaying opposite directional effects in YRI and the European populations in single SNP analysis. For example, the A allele of SNP rs6008600 shows consistently strong negative effect in the four European populations, whereas in YRI the same allele displays a highly significant positive effect (Fig 7). The multi-SNP fine mapping offers a compelling explanation for this phenomenon: two independent *cis*-eQTL signals were identified in the nearby regions, and interestingly, SNP rs6008600 tags one signal in YRI (*r*^2^ ≈ 0.73), whereas in the European populations, it is highly correlated (e.g., in CEU *r*^2^ ≈ 1) with the other signal (Fig 7).

**Fig. 7.**
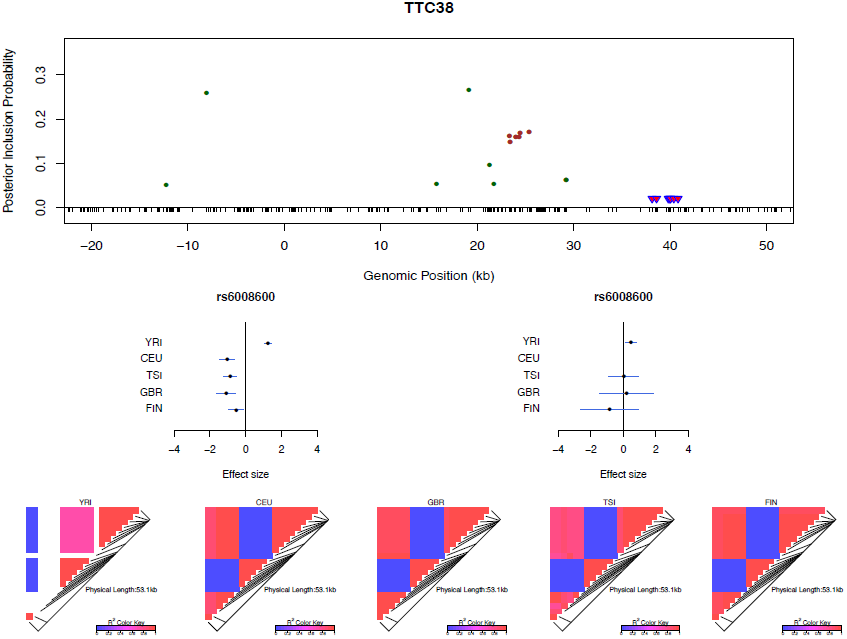
Multi-SNP analysis explains strong effect size heterogeneity observed in single SNP analysis.SNP rs6006800 and the SNPs in LD are labeled by the purple triangles in the top panel. They display strong but opposite effects in the European and YRI populations when analyzed alone. The middle left panel shows the effect sizes of rs6008600 separately estimated in the five populations by the single SNP analysis. The top panel shows the multiple *cis*-eQTL analysis result. SNPs with PIPs ≥ 0.02 are plotted. The result suggests that there are two independent signals in the region, with one represented by SNPs colored in green, and the other represented by SNPs colored in brown. The sums of the green and brown SNPs are both very close to 1. SNP rs6008600 and the SNPs in LD all have PIPs ~ 0 in the multiple *cis*-eQTL analysis. The middle right panel shows the effect sizes of rs6006800 estimated from the multiple linear regression models controlling for the two independent signals in each population: the genetic association observed in the single SNP analysis is seemingly “explained away” by the two independent signals identified by the fine mapping analysis. The bottom panel shows LD heatmaps between SNPs highlighted in the top panel (green, brown SNPs and the SNPs labeled by the purple triangles) in the five populations. Some of the green SNPs are monomorphic in YRI. The opposite effects of rs6006800 are clearly explained by the varying LD patterns: rs6006800 is in high LD with the brown SNPs in YRI, whereas in European populations it tags the green SNPs.

In analyzing the GEUVADIS data, we found the above phenomenon is not uncommon. To further investigate, we followed the procedure described in [21] and quantified the heterogeneity of eQTL effects for all gene-SNP pairs of the identified eGenes in a single SNP analysis. More specifically, for each SNP, we separately computed a Bayes factor (BF_fix_) assuming *constant* eQTL effects across populations (i.e., the fixed effect model) and a Bayes factor (BF_maxH_) assuming completely *independent* eQTL effects across populations. As the most extreme opposite to the fixed effect model, the independent effect assumption corresponds to a model allowing the maximum level of effect size heterogeneity [21]. We then computed a ratio, BF_maxH_/BF_fix_, which itself is a valid Bayes factor measuring the consistency of eQTL effects across population: a large value of the ratio (≫ 1) indicates that the data favor the maximum heterogeneity model and the eQTL effects lack of consistency across populations, whereas a small value (≪ 1) shows that the data support the fixed effect assumption. Based on this analysis, we identified 263 eGenes containing at least one SNP for which BF_maxH_/BF_fix_ ≥ 10^5^, i.e., the evidence for inconsistent eQTL effects is overwhelming in single SNP analysis. Most of those 263 gene-SNP pairs display a pattern of seemingly population-specific effects. For example, the value of BF_maxH_/BF_fix_ for SNP rs6008600 in the above example is ~ 10^24^. For those 263 most extremely heterogeneous single SNP association signals (the list is provided in S1 Dataset), we repeated the calculation of the heterogeneity measure but controlled for the consistent *cis*-eQTL signals identified in our multi-SNP analysis. Fig 8 compares the distributions of the BF_maxH_/BF_fix_ in the single and the multi-SNP analysis models, respectively. The result indicates a striking reduction in heterogeneity by considering multi-SNP analysis models: the median and maximum values of BF_maxH_/BF_fix_ statistics reduce to 1.1 and 10^3^, respectively. This result seemingly suggests that a large proportion of observed heterogeneity in single SNP eQTL mapping is attributed to the varying LD patterns across populations. As a result, when the LD patterns are explicitly considered/modeled in multi-SNP eQTL analysis, we observe more consistent eQTL effects across populations.

**Fig. 8.**
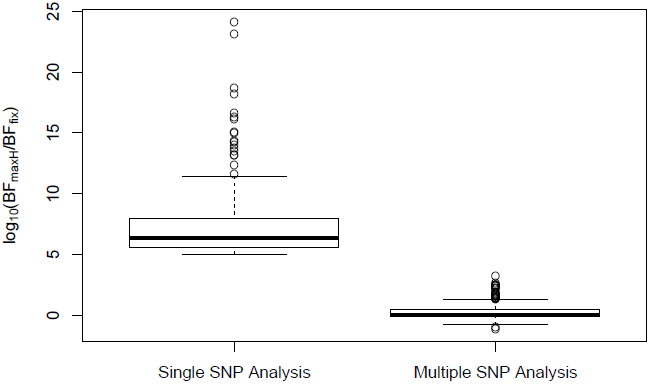
Distributions of SNP effect size heterogeneity from single and multi-SNP analyses. The heterogeneity of SNP effects across the five populations is measured by log_10_[BF_maxH_/BF_fix_], for which large values suggest highly heterogeneous (i.e., inconsistent) genetic effects in different populations. Each point on the plot represents a SNP in a unique gene showing large effect size heterogeneity in the single SNP analysis (log_10_[BF_maxH_/BF_fix_] ≥ 5). When controlling the consistent *cis*-eQTL signals identified by the fine mapping approach in the multi-SNP analysis, we observe a great reduction of effect size heterogeneity.

Although our multi-SNP *cis*-eQTL mapping method confidently identifies independent association signals accounting for LD, in most examples we have examined, it usually cannot pinpoint a single causal SNP by fully resolving LD relying only on the association data. This is because highly correlated SNP genotypes are nearly “interchangeable” in our statistical association models, and therefore not identifiable. As a consequence, there are many combinations of SNPs showing equivalence or near equivalence based on observed data. Reporting a single “best” model while ignoring its intrinsic uncertainty can be highly problematic. In the previously mentioned example of gene *LHPP*, we found that a large proportion of the 134 reported posterior models by the MCMC algorithm exhibit very similar likelihood and posterior probabilities, and the maximum posterior model probability is only 0.02. Our Bayesian fine mapping approach summarizes the measure of uncertainties using the posterior probabilities both at model and SNP levels, which can be naturally carried over to downstream analysis.

#### Functional Annotations of eQTLs

Using the GEUVADIS data and integrating genomic annotations, we set out to investigate the enrichment of certain functional genomic features annotated at SNP level in *cis*-eQTLs. To this end, we conducted the multiple *cis*-eQTL analysis for all 11,858 genes using the prior (5) to obtain PIPs for all gene-SNP pairs, and then examined the correlations between the resulting PIPs and the genomic features of interest. As a side note, notwithstanding the conceptual difference between priors (2) and (5), we noted that the overall results of multiple *cis*-eQTL analysis for identified eGenes are markedly similar.

We first examined the relationship between the abundance of eQTL signals and the SNP distances to the TSS of their respective target genes. For this purpose, we grouped all the *cis*-SNPs into non-overlapping 1Kb bins according to their distances to the respective TSS. We then computed the posterior expected numbers of eQTL signals within each bin by summing over the PIPs of all the *cis*-SNPs falling in the bin. The results are summarized in Fig 9, which displays a strong enrichment of *cis*-eQTL signals nearby the TSS. It is clear that *cis*-eQTLs tend to cluster around the TSS, and the decay of eQTL signals away from the TSS is fast. In particular, Fig 9 suggests that 50% *cis*-eQTL signals are concentrated within 20kb of TSS. Although this phenomenon is well known [5, 18], our results display a much cleaner and more striking pattern comparing to the previous reports. In particular, it is worth noting that, comparing to the similar analysis based on single SNP association test results, our result indicates a much faster rate of decay in abundance of *cis*-eQTLs away from the TSS. This is most likely because we considered multiple *cis*-eQTL signals for a target gene, and our use of PIP accounted for the uncertainty of eQTL calls.

**Fig. 9.**
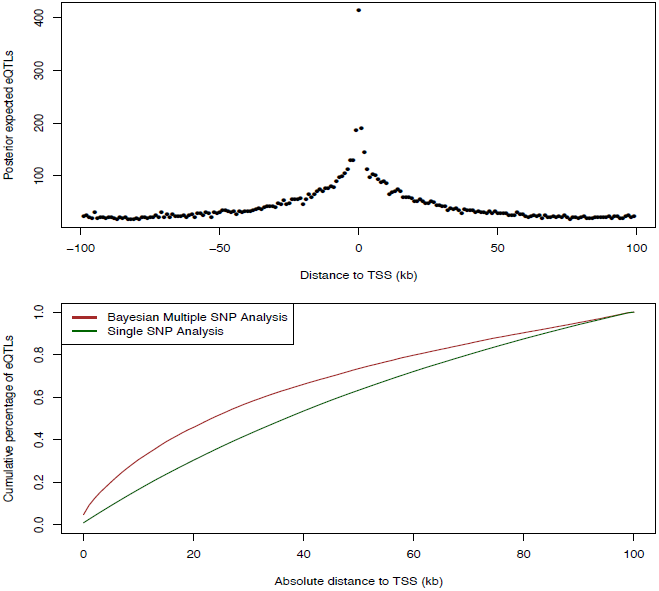
Enrichment of *cis*-eQTLs with repect to distance to TSS. The top panel shows the distribution of posterior expected number of *cis*-eQTLs with respect to SNP distance to TSS. The bottom panel shows the estimated cumulative percentage of *cis*-eQTLs with respect to SNP disance to TSS from two different methods. The brown line represents the estimate using the PIPs from the proposed Bayesian approach. The dark green line represents the estimate from a standard method based on single SNP association results. More specifically, the dark green line is estimated by counting the number of SNPs exceeds FDR 5% threhold in each distance bin. The brown line shows a much faster decay in abundance of *cis*-eQTL signals away from TSS.

We next asked whether eQTLs are enriched for genetic variants disrupting TF binding. Answering this question helps reveal the underlying regulatory logic linking between transcription factor binding and gene expression [17, 18, 28, 29]. To this end, we used the base-pair resolution quantitative annotation of binding variants derived from DNase-seq data for LCLs in the ENCODE project (the detailed description of this annotation and the methods used to generate it can be found in [30] and also in S1 Text). Briefly, for each TF motif a re-calibrated CENTIPEDE model [11] is built to integrate its DNase-seq footprint shape characteristics and the TF sequence preference of binding. Then, each genetic variant from the 1000 Genomes project is annotated if it overlaps with a CENTIPEDE footprint for a TF. The sequence model for the TF is also used to evaluate the prior probability for each of the two alleles to predict the impact of the genetic variant on binding. There are three mutually exclusive scenarios using this annotation:

- SNPs that are not located in any DNase I footprint region, or in brief, *baseline SNPs*
- SNPs that are in a footprint region but predicted to have little or no impact on TF binding based on the sequence model, or in brief, *footprint SNPs*
- SNPs that are in a footprint region and predicted to strongly affect TF binding, or in brief, *binding variants*

About 4% of the total 6.7 million interrogated *cis*-SNPs are annotated as binding variants, and another 4% are annotated as footprint SNPs. Here, we hypothesized that binding variants have a higher enrichment level than footprint SNPs in eQTLs. To this end, we examined the association between the SNP-level binding annotation and PIPs for *cis*-eQTLs. To control for the fact that footprints are also enriched in the close neighborhood of the TSS of transcribed genes, we additionally controlled for SNP positions with respect to the TSS in the analysis.

We provide the complete inference results, including the estimates of the enrichment parameters for all bins defined by the distance to the TSS, in S2 Dataset. In brief, our main findings from this analysis are:

- binding variants are 1.49 fold (with 95% confidence interval [1.38, 1.63]) more likely than baseline SNPs to be eQTLs, their enrichment in eQTLs is statistically highly significant (*p*-value = 4.93 × 10^−22^)
- footprint SNPs are 1.15 fold (with 95% confidence interval [1.04, 1.27]) enriched in eQTLs, comparing to baseline SNPs, with the corresponding *p*-value = 0.0035

This result implies that the sequence variants that potentially disrupt TF binding are more likely to have a functional impact on gene expression. In comparison, footprint SNPs (those predicted to not affect binding) are less likely to affect expression, and this is reflected with a relatively lower level of enrichment significance for eQTLs.

Finally, we returned to the fine mapping analysis and quantitatively incorporated the binding annotation information into the *cis*-eQTL discovery. More specifically, we plugged in the point estimates of the enrichment parameters from the association test to re-compute the Pr(*γ*_*j*_) using the logistic model (6) for each gene-SNP pair, and re-ran the MCMC algorithm with the updated priors. As a result, the priors for binding variants are up-weighted compared with the nearby baseline and footprint SNPs. Because the computation of likelihood function is intact, the PIPs for binding variants are accordingly up-weighted. Using this approach, it becomes plausible to quantitatively distinguish SNPs in perfect LD but belonging to different annotation categories. Take Gene *LY86* (Ensembl ID: ENSG00000112799) as an example (Fig 10). Before incorporating any annotation information, each *cis*-SNP was equally weighted *a priori* and the multiple *cis*-eQTL analysis identified three independent signals, two of which are represented by clusters of highly correlated SNPs. Utilizing the quantitative annotation information, the fine mapping procedure still confidently identifies the three original eQTL signals, however two binding variants become the top associated SNPs (according to the PIPs) in each respective cluster of SNPs that are indistinguishable from several other SNPs in the original analysis (Fig 10).

**Fig. 10.**
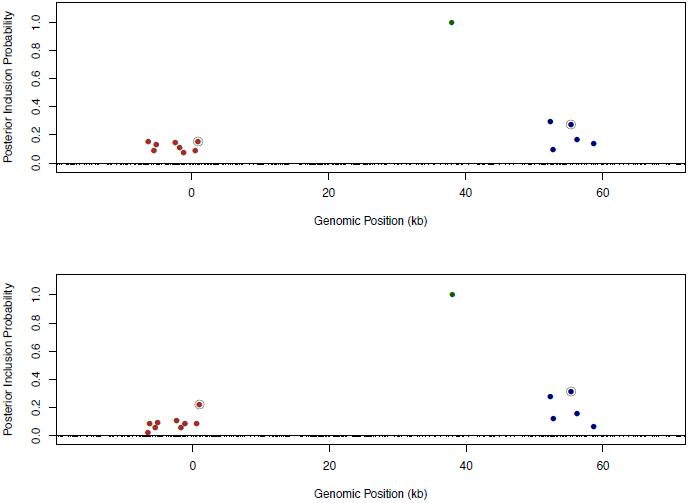
Comparison of fine mapping results of gene *LY86* before and after incorporating functional annotations. The top panel is based on the analysis using prior (5) without annotation information. The bottom panel is based on the analysis using prior (6) incorporating SNP distances to TSS and the annotations for binding variants and footprint SNPs. In both panels, SNPs with PIPs ≥ 0.02 are plotted. There are clearly three independent *cis*-eQTLs in the region, represented by different colors of SNPs. SNPs in smae colors are in high LD. The sums of PIPs from SNPs in same colors are all close to 1. The circled points are predicted binding variants. It is clear that binding variants are up-weighted when annotation information is incorporated into the fine mapping analysis.

In the set of 526 genes that we confidently identified as harboring only one single *cis*-eQTL signal, when applying the equal prior without any annotation information, we found that binding variants and footprint variants are top associated SNPs in 11% and 8% of the genes (or 60 and 43 genes in number), respectively. Using the priors incorporating quantitative binding annotation and SNP distance to the TSS, the percentages of genes with binding variants and footprint SNPs as top associated SNPs increase to 16% and 11% (or 85 and 57 genes in number), respectively.

## Discussion

We have presented an integrative analysis framework to perform fine mapping analysis jointly from cross-population samples while incorporating functional annotations. Our core statistical methods are built on and naturally extend the established Bayesian framework for association analysis of genetic data from heterogeneous groups [14, 15, 21]. The most notable features of our statistical framework are its ability of multi-SNP analysis and integration of SNP-level genomic features. Through simulation studies, we have demonstrated the power and efficiency of the proposed approaches. For the first time, we have applied this framework for multiple *cis*-eQTL fine mapping analysis in a cross-population meta-analytic setting using the GEUVADIS data.

Our analysis reveals that with an aggregated sample size of around 400, multiple independent *cis*-eQTL signals can be confidently identified in many genes, which makes evident the necessity of considering multiple association signals in eQTL studies. The commonly applied strategy for mapping additional association signals is to conduct conditional analysis, which can be viewed as a step-wise variable selection algorithm. Besides lack of power, one additional disadvantage of conditional analysis is that it only reports a single “best” model in the end, and completely ignores its uncertainty. From many of our examples shown in this paper, it is clear that in most cases there is typically a great deal of uncertainty in determining the truly causal SNPs due to LD, and the posterior probabilities of the “best” models are often unimpressive. Consequently, it is generally inappropriate to solely rely on the information from the “best” model in the downstream analysis (as we have illustrated in the power simulations of enrichment testing). In comparison, in addition to gaining power in identifying multiple independent *cis*-eQTL signals, our Bayesian approach provides much more comprehensive information that fully conveys the uncertainty of the inference result, and the quantified uncertainty information is naturally propagated in our integrative analysis of genomic features.

In this paper, we have employed a two-step procedure that first screens eGenes by performing gene-level hypothesis testing, and then carries out multi-SNP analysis for the identified eGenes. This procedure is analogous to the fine mapping procedure that is commonly used in GWAS, where interesting loci are ranked and selected by single SNP association testing before an in-depth analysis focusing on each flagged high priority locus. We find this procedure not only yields considerable computational savings, but also provides a sound argument to specify the prior inclusion probability Pr(*γ*_*j*_).

It should be noted that our proposed analysis procedures are completely applicable for fine mapping of general QTLs in either a meta-analytic or single study setting. They can be further extended to the applications of fine mapping analysis where subgroups of eQTL data are formed by different tissues [4, 14] or cellular conditions [6, 8]. Comparing to the meta-analytic setting considered in this paper, the parameter space of {*γ*_*j*_} in those applications is more complicated (which includes 2^*s*^ potential values, where *s* is the number of subgroups/tissues). Nevertheless, [14] has provided a principled way to “learn” the priors on possible values that *γ*_*j*_ can take by pooling information across genes through a hierarchical model.

Our fine mapping results of eQTLs also demonstrate the benefit of utilizing cross-population samples in genetic association studies. Most importantly, the population heterogeneity of local LD patterns serves as an efficient filter that narrows down the regions harboring casual eQTLs. Nevertheless, varying LD patterns can cause some SNPs to display large degree of heterogeneity across populations in their estimated effect sizes from *single SNP analysis*: in the extreme cases, a SNP may appear to possess strong “population specific” effects. As we acknowledge that genuine population specific eQTLs are certainly interesting phenomena and very much likely exist, we suggest *interpreting* highly heterogeneous eQTL signals from single SNP analysis with caution. In our view, it may be necessary to carry out multi-SNP analysis, as we have demonstrated in this paper, to simply rule out the possibility that the seemingly population specific effects are artifacts due to varying LD patterns.

Our Bayesian inference framework naturally incorporates functional annotations in fine mapping eQTLs across population groups. This feature allows us to quantitatively assess the enrichment of certain functional features in eQTLs, and in turn to use the quantified enrichment information to prioritize annotated SNPs for fine mapping analysis. Overall, our model for integrative eQTL mapping analysis is similar to those presented in [17, 18]. However, these previous approaches make simplifying assumptions to restrict at most one *cis*-eQTL per gene, such that single SNP association results can be directly used. Our method relaxes this assumption and is fully integrated into our multi-SNP analysis procedure. In addition, our use of CENTIPEDE annotation to examine the relative enrichment of binding variants and footprint SNPs is also novel. Although it is largely expected that binding variants are enriched in eQTLs, it is important to note that the level of enrichment for footprint SNPs is much lower than that for binding variants. Interestingly, this finding seems concurring with the results reported by [30] where the relative enrichment of binding variants vs. footprint SNPs in other cellular and organismal phenotype QTLs is examined.

Last but not least, our multi-SNP fine mapping analysis of the GEUVADIS data has created a comprehensive resource for the community to gain better understanding of the genetic basis of gene regulation. With proper uncertainty assessments, our results enable follow-up experimental validations and functional studies of causal genetic variants that alter gene regulation. They also provide a unique and powerful resource to study population genetics of expression traits. For example, we found that the distribution of *F*_*st*_ of eQTL signals consistent across populations has a shorter tail than the distributions of random SNPs (Kolmogorov-Smirnov test *p*-value = 0.001). Nevertheless, when examining the *F*_*st*_ distributions between primary and secondary *cis*-eQTL signals (stratified by the association strength in single SNP analysis with respect to their target genes), we did not find statistical evidence to differentiate the two (Kolmogorov-Smirnov test *p*-value = 0.77). There are many interesting aspects to be explored with our analysis results, and we will leave the more in-depth investigations for our future work.

## Materials and Methods

The computational methods used in the analysis are implemented in the software packages FM-eQTL (manuscript in preparation) and eQTLBMA [14]. They are freely available at https://github.com/timflutre/eqtlbma and https://github.com/xqwen/fmeqtl

The fine mapping results of all 6,555 identified eGenes, including the scatter plots of PIPs, forest plots of top models, and the detailed summaries from separate linear regression analysis are made available on the website http://www-personal.umich.edu/~xwen/geuvadis/

### Data Pre-processing

The genotype and RNA-seq data were downloaded from the GEUVADIS project website. We selected 420 samples whose genotypes are directly measured in the 1000 Genomes project. The samples are evenly distributed in the five population groups, and the detailed breakdown of samples by population is shown in Table 2.

For the RNA-seq data, we used a slightly more stringent threshold than the original analysis [20] to select genes that are expressed in all five populations. Specifically, for each selected gene, we required > 90% individuals in each population group to have RPKM ≥ 0.1. From the 17,361 Ensembl genes that passed this filter, following the original analysis, we selected a subset of 11,838 genes consisting of annotated protein-coding genes and lincRNAs according to GENCODE [31] release 17. We log transformed the RPKM values, and used the pipeline employed in [29, 32] to remove the effect of GC content on expression measurements. We then followed the same strategy as described in [20] to remove latent confounding factors using the software package PEER [33]. However, unlike the original analysis, we ran PEER for each population group separately. In the end, we removed 15, 13, 15, 20 and 20 PEER factors for samples from YRI, CEU, TSI, GBR and FIN, respectively. Finally, the expression levels of each gene were quantile normalized across individuals separately in each population group.

For genotype data, we filtered out SNPs whose sample allele frequencies < 0.03 in the *overall* samples across population groups. Note, we did not apply the allele frequency filter in each population group. The SNPs passing this filter must have sample allele frequencies ≥ 0.03 in at least one population group. In general, as discussed in the Results section, rare SNPs do not impose any statistical or computational problems for our analysis. Nevertheless, removing SNPs that are not likely informative in *any* population group helps improve computational efficiency. Following [17, 32], we defined a 200kb *cis* region for each gene centered at its TSS. In total, the final data set contains 6.7 million gene-SNP pairs.

### Gene-level Analysis of *cis*-eQTLs

For each gene, we tested the null hypothesis that asserts no *cis*-eQTLs. Specifically, we adopted the Bayesian hypothesis testing procedure discussed in [14]. Essentially, [14] assumes a Bayesian model that is mostly similar to (1), except for an additional simplifying assumption, “at most one *cis*-eQTL per gene”. Given a gene with *p cis*-SNPs, this additional assumption reduces the possible alternative scenarios into *p* single SNP association models, for which a gene-level Bayes factor can be easily computed analytically by Bayesian model averaging. In the context of eQTL mapping in multiple tissues, [14] considered all possible configurations, i.e., *γ*_*j*_ values, for each assumed associated SNP, whereas in our analysis of the GEUVADIS data we only allowed *γ*_*j*_ ∈ {**0**, **1**} to focus on discovering population consistent *cis*-eQTLs.

Briefly, for the alternative model where the *j*-th SNP is assumed the lone eQTL, the linear model (1) is simplified to

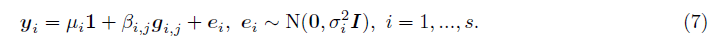

We modeled the correlation of genetic effects, *β*_*i, j*_’s across population groups through the following prior specification,

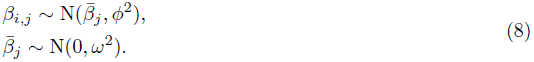

Equivalently, the above prior can be represented by a multivariate normal distribution by integrating out the average effect parameter 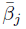, i.e.,

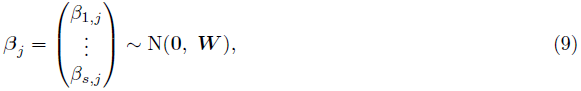

where the *s* × *s* variance-covariance matrix ***W*** is given by

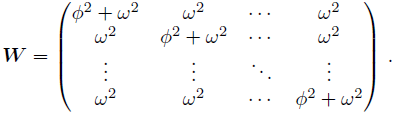

The parameter *ϕ*^2^ + *ω*^2^ characterizes the overall genetic effects of SNP *j*, and *ϕ*^2^/(*ϕ*^2^ + *ω*^2^) represents the degree of heterogeneity across population groups. Following [14, 15] we considered the values of *ϕ*^2^ + *ω*^2^ from a set *E* = {*ϕ*^2^ + *ω*^2^: 0.1^2^, 0.2^2^, 0.4^2^, 0.8^2^, 1.6^2^} which covers a wide range of plausible magnitude of genetic effects. We allowed limited degree of heterogeneity by taking *ϕ*^2^/(*ϕ*^2^ + *ω*^2^) values from the set *H* = {*ϕ*^2^/(*ϕ*^2^ + *ω*^2^): 0, 0.1} which reflects our prior belief that effects of genuine eQTL signals should be highly consistent across population groups. Overall, we considered a combination of |*E*| × |*H*| grid for (*ϕ*^2^, *ω*^2^) values for each alternative model. Given this model, a single SNP Bayes factor, BF_*j*_, can be analytically evaluated following [14, 15], and the corresponding gene-level Bayes factor is obtained by Bayesian model averaging as

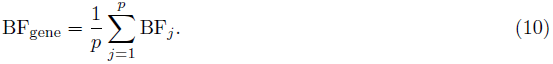

Upon obtaining the gene-level Bayes factors, we used the methods implemented in the software package eQTLBMA [14] to select eGenes at FDR 5% levels. eQTLBMA implements two types of FDR control procedures: one is a permutation based procedure which converts a gene-level Bayes factor to a corresponding *p*-value and control FDR using Storey’s procedure; the other procedure is based on the EBF procedure discussed in [34] which directly works with gene-level Bayes factors and avoids any permutations. We found that the latter approach is much more computationally efficient, however slightly conservative. The results reported in the Results section are based on the EBF procedure.

The power gain of the Bayesian gene-level testing procedure in identifying consistent association signals across multiple populations/subgroups and its comparison with commonly applied frequentist approaches have been fully demonstrated in [14] and [21] through simulations and real data analysis. Here, we examined a few known factors that may potentially affect the power of the gene-level analysis in the GUEVADIS data.

Using the hierarchical model proposed in [14], we performed the analysis described in [21] to estimate the effect size heterogeneity of eQTL signals across populations: our choice of the heterogeneity parameters is mostly informed by this analysis. More specially, we used a full spectrum of grid values for the heterogeneity parameter *ϕ*^2^/(*ϕ*^2^+*ω*^2^) from a comprehensive set *H* = {*ϕ*^2^/(*ϕ*^2^+*ω*^2^): 0.0, 0.1, 0.2,…, 0.9, 1.0}. We then estimated the probability weight on each grid value by pooling information across all genes using the EM algorithm implemented in eQTLBMA. The estimates show that the majority of the probability mass is concentrated in very low heterogeneity levels (Table 3), and the mean heterogeneity E[*ϕ*^2^/(*ϕ*^2^ + *ω*^2^) | ***Y, G***] = 0.060, i.e., on average, the eQTL effects are very consistent across populations. This result is also highly concordant to the findings in [21] where eQTL data from microarray experiments across European, African and Asian populations were examined, and further justifies our selection of heterogeneity parameter values in the analysis.

**Tab. 3.** Estimated heterogeneity levels of eQTL effects across populations

| Heterogeneity Level ($\frac{\phi^2}{\phi^2 + \omega^2}$) | Estimated Weight |
| --- | --- |
| 0.0 | 0.651 |
| 0.1 | 0.204 |
| 0.2 | 0.084 |
| 0.3 | 0.035 |
| 0.4 | 0.014 |
| 0.5 | 0.006 |
| 0.6 | 0.003 |
| 0.7 | 0.002 |
| 0.8 | 0.000 |
| 0.9 | 0.000 |
| 1.0 | 0.000 |

The estimates are obtained from the hierarchical model described in [14, 21]. For each grid value of 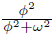, ranging from 0.0 to 1.0, we estimate its probability weight by pooling information of all gene-SNP pairs. The grid value 0 indicates a fixed (i.e., the most consistent) eQTL effect across all populations and has the most estimated weight. The grid value 1 indicates completely independent eQTL effects across populations. The results indicate that across population groups, on average, the eQTL effects exhibit low level of heterogeneity.

Among other factors impacting eGene discoveries, we noted that the number of *cis*-SNPs is negatively correlated with the gene-level local false discovery rate (lfdr). That is, on average, genes harboring more *cis*-SNPs tend to exhibit stronger gene-level association signals. The Spearman’s rank correlation between the two quantities is modest (-0.20), nonetheless statistically highly significant (*p*-value < 2.2 × 10^−16^).

### Multi-SNP Analysis of *cis*-eQTLs

In our multi-SNP analysis, we no longer assumed “one *cis*-eQTL per gene”, and considered the full range of alternative scenarios described by model (1). To make joint inference with respect to **Γ** = {*γ*_*i*_,…, *γ*_*p*_}, we further specified effect size distribution,

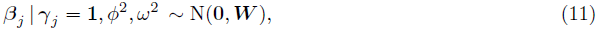

where ***W*** is constructed in the same way as in the gene-level analysis. In particular, we used the same set of grid values of (*ϕ*^2^, *ω*^2^) in the gene-level analysis. Unconditional on *γ*_*j*_, the prior on *β*_*j*_ is a type of “spike-and-slab”, where the “slab” is represented by a mixture of multivariate normal distribution, and the vast majority of the prior probability mass, 1 − 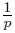, is assigned to the “spike” (i.e., a point mass at 0).

The described linear model system including the prior specification (i.e., SSLR), is a special case of the general system considered by [15]. Given a specified **Γ** value, a Bayes factor, BF(**Γ**), contrasting to the trivial null model, **Γ** ≡ **0**, can be analytically approximated by applying the result discussed therein. With this result, the posterior probability, Pr(**Γ** | ***Y, G***), can be computed up to an unknown normalizing constant, i.e., Pr(**Γ** | ***Y, G***) ∝ Pr(**Γ**) BF(**Γ**),. We implemented a Metropolis-Hastings algorithm, similar to what is discussed in [15] for multivariate linear regression model (MVLR), to efficiently traverse the space of 2^*p*^ possible **Γ** values. In particular, we designed a novel proposal distribution that utilizes marginal and conditional analysis results to prioritize SNPs with strong marginal or conditional association signals. In practice, we observed that the resulting MCMC algorithm achieves fast mixing. The details of the algorithm is provided in S1 Text (section S.1). In the end, the MCMC algorithm yields a set of **Γ** samples from the posterior distribution, from which we computed the PIP for each SNP by marginalization.

The posterior expected number of independent *cis*-eQTLs and its variance for each gene are obtained by computing the sample mean and variance of the number of non-zero *γ*_*j*_’s in each posterior model. Equivalently, the posterior expected number of *cis*-eQTLs can be computed by the sum of PIPs, i.e.,

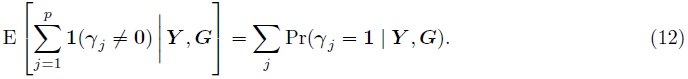

For the fine mapping analysis of the GEUVADIS data, we applied the MCMC algorithm for each identified eGene individually. We carried out 25,000 burnin steps and 50,000 repeats for each MCMC run. Taking advantage of parallel processing, we performed multiple-SNP analysis for multiple genes simultaneously in a distributed computing environment, which greatly reduced the overall computational time. The MCMC run for the eGene with the most *cis*-SNPs, *HLA-DRB1* (Ensembl ID: ENSG00000196126, with 11,400 *cis*-SNPs), took 30 minutes on a computer with a single Intel Xeon 2.13GHz 8-core CPU. For average eGenes with ~ 2,000 *cis*-SNPs the running time is approximately 3 to 4 minutes on the same machine. We ran our fine mapping analysis for GEUVADIS data with 8 parallel threads, the overall computation took about 30 hours.

Although the MCMC output conveys the full information of the fine mapping results, parsing the information manually to identify independent eQTL signal clusters can still be non-trivial. To tackle this problem, we designed and implemented a simple hierarchical clustering based algorithm that automatically parses the MCMC output and aides identifying independent eQTL signal clusters. The details of the algorithm are described in the S1 Text (section S.4).

#### Simulation Study

We performed numerical simulations to evaluate the performance of the proposed Bayesian multi-SNP analysis approach. For this purpose, we simulated the expression levels of 1,500 genes. The genotypes of *cis*-SNPs in each gene were assembled using the real genotype data from the GEUVADIS project. More specifically, we randomly selected 100 genomic regions each containing 25 consecutive SNPs, and directly took the observed genotypes of those 2,500 SNPs from each GEUVADIS population. Consequently, there are generally modest to high levels of LD within each region, and unremarkable levels of LD between regions. The resulting genotype data also maintain the variations in genotype frequency and local LD patterns (within each 25-SNP region) across populations. For each gene, we randomly assigned 1 to 4 population-consistent eQTL SNPs and generate the expression levels for all five populations using a system of linear models (details are described in S.5 of S1 Text). We repeated this procedure to generate the genotype-expression data for all 1,500 genes.

We analyzed the simulated data set using the following three different *cis*-eQTL mapping approaches:

1. the proposed Bayesian multi-SNP mapping method
2. a fixed effect single SNP meta-analysis approach
3. a conditional analysis approach (i.e., a forward step-wise selection procedure) built upon the single SNP meta-analysis procedure in 2

We evaluated the three different mapping methods by their abilities in correctly identifying the (25-SNP) regions harboring the true causal variants. For the Bayesian approach, we computed a regional PIP by summing over the SNP-level PIPs of each region and selected the regions whose regional PIPs exceed a pre-defined probability threshold. For the single SNP analysis, we selected regions whose minimum *p*-values of the including SNPs are below a pre-defined *p*-value threshold. Finally, we performed the conditional analysis by a forward step-wise selection procedure using a pre-fixed *p*-value threshold. For both conditional and single SNP analyses, the SNP-level *p*-value was computed from the fixed effect meta-analysis model (the details are provided in S.5 of S1 Text). For each method in comparison, we varied the corresponding threshold values in a range from very stringent (i.e., few false positives) to very relaxed (i.e., few false negatives), and for each threshold value we recorded the corresponding true positives and false positives from the identified regions.

Fig 1 shows the comparison of the three approaches based on the simulated data set. It is clear that multi-SNP analysis methods (both the Bayesian and conditional analysis approaches) outperform single SNP analysis when multiple *cis*-eQTL signals are commonly present. More importantly, for any given fixed false positive value, the proposed Bayesian method always yields the most true positives, outperforming the conditional analysis approach. We further investigated the power gain in Bayesian multi-SNP analysis by stratifying the simulated genes by their number of *cis*-eQTLs. The results, shown in Fig 11, indicate that when a gene contains only a single *cis*-eQTL, all three methods have very similar power. However, when multiple *cis*-eQTLs are harbored by a single gene, the Bayesian multi-SNP approach shows consistently superior power over the two alternative procedures.

**Fig. 11.**
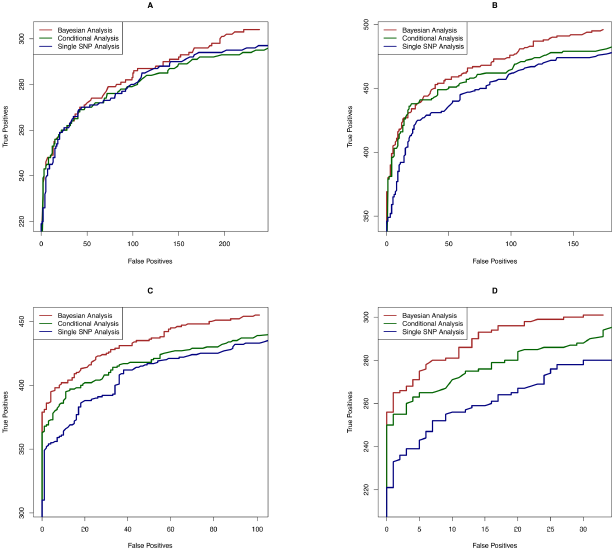
Power comparison in identifying causal eQTL regions stratified by the number of true eQTL Signals in each gene. Panels A, B, C and D show simulation results for genes containing 1, 2, 3 and 4 *cis*-eQTLs, respectively. In each panel, three competing methods are compared: the proposed Bayesian multi-SNP analysis method (brown line), a conditional meta-analysis method (dark green line) and a single SNP meta-analysis approach (navy blue line). Each plotted point on each panel represents the number of true positive findings versus the number of false positive findings of a given method at a particular threshold. In the simulation study, the numbers of genes containing 1, 2, 3 and 4 *cis*-eQTLs are 579, 457, 307 and 157, respectively. For genes that harbor only a single eQTL, all three methods yield similar performance. However, when there exist more eQTLs in a gene, the proposed Bayesian multi-SNP approach shows superior power over the two competing procedures.

Additionally, we examined the calibration of the reported regional PIPs using the simulated data set (Fig 13). The results indicate that the reported regional PIPs are generally well calibrated, i.e., the reported probability values truly reflect the frequencies of the true causal regions over many independent observations. We observed that the small to modest PIP values tend to be slightly conservative, which may be explained by the use of prior model (2). This calibration property ensures a natural Bayesian FDR control [35] in determining the PIP threshold for selecting the causal regions. Fig 12 plots the comparison between estimated FDR from PIPs and the true realized FDR in the simulated data. The two quantities are in very good agreement. For example, in our simulation at the estimated 5% FDR level, we select 1432 causal regions (with PIP cutoff = 0.424) of which the realized true false discovery percentage is 4.6%. In comparison, we found it difficult to determine an appropriate *p*-value cutoff for conditional and single SNP analyses. The commonly applied Bonferroni correction threshold seems too stringent in our simulation setting: the single SNP analysis and the conditional analysis select only 1070 and 1113 regions (with 2 and 1 false positives), respectively.

**Fig. 12.**
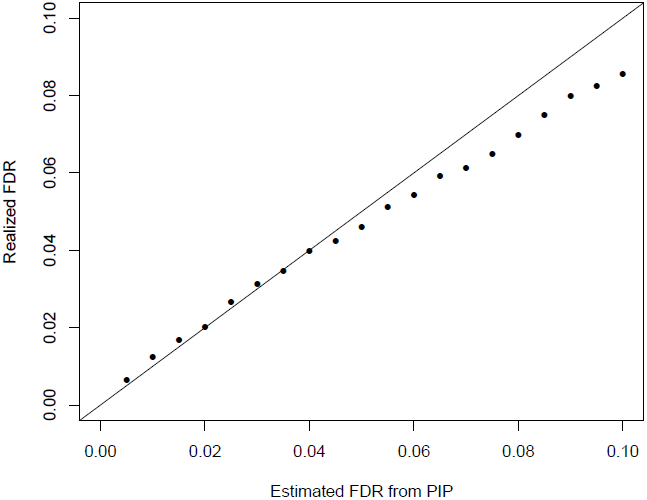
The comparison between the realized falase discovery rate and the estimated false discovery rate using the PIPs from the proposed Bayesian method in the simulation study. The plot indicates that the FDRs estimated by PIPs are mostly accurate and become slightly conservative for large FDR values.

**Fig. 13.**
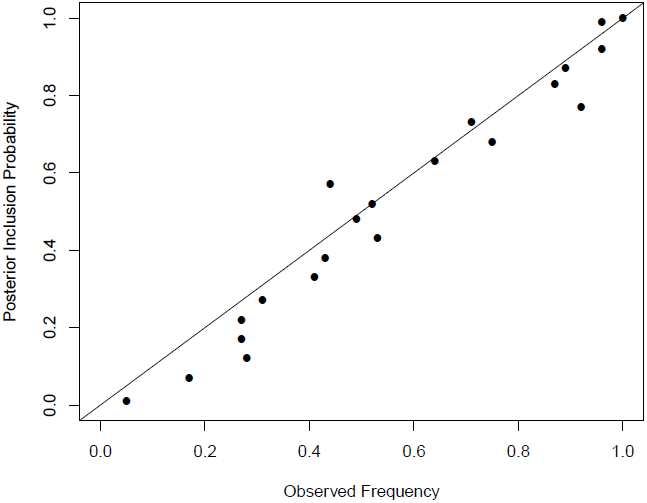
Calibration of PIPs from multiple *cis*-eQTL analysis. Using the simulated data set, we grouped the reported 1500 × 100 regional PIP values, ranging from 0.0 to 1.0, into 20 bins of width 0.05. For each bin, we plot the mean PIP value and the corresponding frequency of true causal regions represented in the bin. All points are closely located to the diagonal line, indicating that the reported PIPs are relatively well-calibrated.

### Enrichment Testing of Genomic Annotations

To test the enrichment of a particular genomic annotation in eQTL signals, we examine the associations between SNP level annotations of interest and the PIPs from our multi-SNP fine mapping approach computed under the null hypothesis of no enrichment.

Without loss of generality, we consider a single annotation and use 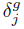 to denote the annotated value of SNP *j* in gene *g* (the additional super-script for gene emphasizes the annotation is specific to each gene-SNP pair). Our testing procedure starts by computing PIPs for all gene-SNP pairs using the proposed multi-SNP *cis*-eQTL mapping method with the exchangeable prior specification (4). Under the null hypothesis, the resulting PIPs should be independent of the annotation information. We then fit a logistic regression model

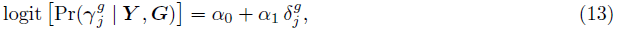

and perform the Wald test with respect to *α*_1_.

The commonly applied enrichment testing approach typically classifies the latent association status of each SNP 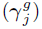 into 0 or 1 depending on the single SNP association test results (usually *p*-values), and then test their associations with the annotations of interest. In comparison, the use of PIPs in our proposed method presents at least two obvious advantages: first, it naturally accounts for potentially multiple association signals within a single gene; and second, it fully accounts for the uncertainty in determining 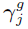 which is ignored by the binary classification approach. These advantages are reflected by the gain of powers in enrichment testing.

#### Simulation Study

We performed simulation studies to evaluate the performance of the proposed enrichment testing procedure.

We selected 2,500 genes from the GEUVADIS data. To ease computation, we only kept 50 randomly selected *cis*-SNPs for each gene. We randomly annotated 20% of the SNPs with a binary feature. For each SNP *j* in gene *g*, we determined its true association status by an independent Bernoulli 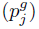 trial, where

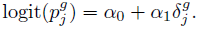

In particular, we set *α*_0_ = −4.59, which implies about 1% of unannotated SNPs are expected to be true eQTLs. We selected values of *α*_1_ from the set {0.00, 0.25, 0.50, 0.75} for different simulations. Once the causal SNPs were determined, we simulated their effect sizes and expression levels of target genes using the same approach described in the simulation study of fine mapping analysis (section S.5 of S1 Text). For each unique *α*_1_ value, we generated 1,000 independent data sets.

We analyzed each simulated data set using the proposed approach. For comparison, we also applied a standard enrichment analysis procedure principally similar to what described in [20, 36]. More specifically, we performed single SNP association tests for all gene-SNP pairs in a simulated data set, then classified the top associated SNPs of each gene as the “causal” eQTLs if their *p*-values pass the significance threshold at FDR 5% level. Finally, we performed a Fisher’s exact test to assess the correlation between the the classified association status and the annotation. S2 Table shows the comparison of type I errors and powers for the two approaches. The type I errors are well controlled for both methods. The proposed enrichment testing approach utilizing PIPs shows much elevated statistical powers in our simulation settings.

### Integrative Fine Mapping Analysis of *cis*-eQTLs

To incorporate genomic annotations into the fine mapping analysis of *cis*-eQTLs, we specify the prior distribution of Pr(*γ*_*j*_) by the parametric function (6), whereas the other parts of the Bayesian multi-SNP analysis model remain intact. For the *g*-th gene and its *j*-th SNP, we re-write (6) using an equivalent vector form as follows

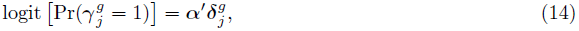

where the enrichment parameter ***α*** is assumed to be shared across all gene-SNP pairs.

Let ***D***^*g*^ denote the collection of the annotation data for gene *g*. For a total of *q* genes, it follows from the Bayes rule that

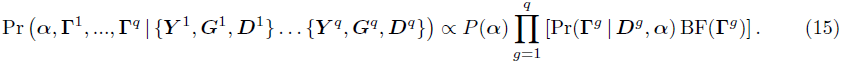

Given a prior specification *P*(***α***) (e.g. a flat prior), the quantities on the right hand side are individually straightforward to compute analytically. It is conceptually easy to modify our MCMC algorithm for the multi-SNP analysis to jointly sample (***α***, **Γ**^1^,…, **Γ**^*q*^). Nevertheless, due to the extremely high dimensionality of the target space (which is approximately the number of gene-SNP pairs, in the case of the GEUVADIS data the number is ~ 6.7 million), the convergence of the MCMC within a reasonable time frame may be in doubt. Furthermore, the computational resources, especially the memory usage, demanded by the MCMC algorithm may be too high to afford in a practical setting.

Alternatively, we consider an empirical Bayes approach which employs an EM algorithm to find the MLE of ***α*** by treating (**Γ**^1^,…, **Γ**^*q*^) as missing data, and perform a final round of multi-SNP analysis (possibly only for eGenes) conditional on the resulting MLE 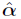. The derivation of the EM algorithm is mostly straightforward, we give the relevant details in S1 Text (section S.2). Briefly at (*t* + 1)-th iteration, in the Expectation step (E-step), we evaluate 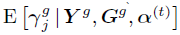 for each gene-SNP pair given the current estimate ***α***^(*t*)^. It should be noted that this conditional expectation is exactly the PIP of each gene-SNP pair which can be obtained by running the multi-SNP analysis for each gene *separately* given the hyperparameter ***α***^(*t*)^. In the Maximization step (M-step), we find *α*^(*t*+1)^ by fitting a logistic regression model relating PIP of each gene-SNP pair to the genomic annotations of the corresponding SNP. Overall, we describe the complete algorithm as an “MCMC-within-EM” algorithm, which is initiated at some arbitrary ***α***^(0)^ value, and iteratively performs multiple *cis*-eQTL mapping using the MCMC algorithm and maximization by fitting logistic regression models until convergence. The main computational benefit of the “MCMC-within-EM” algorithm is that the E-steps involving MCMC runs can be processed in parallel on a distributed computing system, hence the memory requirement is much relaxed.

To illustrate the procedure for estimating ***α***, we simulated a set of eQTL data using the same scheme in the enrichment testing. In particular, we used a single binary annotation and set *α*_1_ = 1.50. We ran the MCMC-within-EM algorithm on this simulated data set with the starting point *α*_1_ = 0, and plotted the estimated *α*_1_ values in each iteration in Fig 14. The figure indicates that the estimates of *α*_1_ quickly converge to the close neighborhood of the true value after 6 to 7 iterations.

**Fig. 14.**
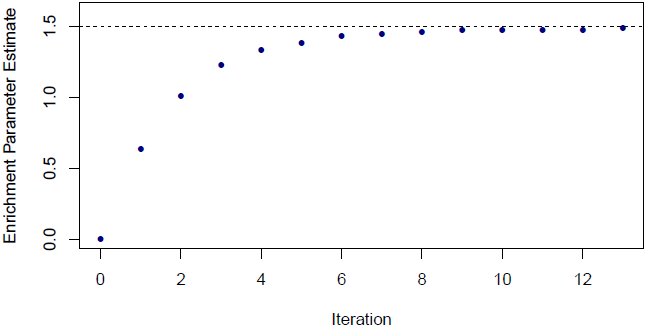
An illustration of the convergence of the enrichment parameter estimates in the proposed MCMC-within-EM algorithm. In this simulated data set, the true enrichment parameter is set to 1.50 (the dotted horizontal line). The algorithm initiate the parameter value at 0 and quickly reaches to the close neighborhood of the true value.

To reduce computation, in practice we analyzed the GEUVADIS data using only a single iteration of EM where all enrichment parameters are initialized at 0. As a consequence, our reported point estimates of the enrichment parameters are likely to be lower bounds of the true values. Nevertheless, as we have shown in the analysis of the GEUVADIS data, this simplified version of the algorithm still improves the resolutions of the fine mapping analysis by up-weighting the SNPs with relevant annotations.

## Acknowledgments

We thank Matthew Stephens, Timothée Flutre, Andrei Stefanescu, Tuuli Lappalainen, Casey Brown and the two other anonymous reviewers for helpful discussions and comments.

## Supplementary Text

### S.1 MCMC Algorithm for Mapping Multiple *cis*-eQTLs

We implemented a Metropolis-Hastings algorithm to perform posterior sampling based on equation (2.3) in the main text. The algorithm is mostly straightforward. To help the Markov chain achieve fast mixing, we implemented a novel proposal distribution based on the result of conditional analysis of multiple *cis*-eQTLs.

We propose two types of simple “local” moves in the MCMC simulations:

1. Change a *γ*_*j*_ value for SNP *j*
2. Swap the values of *γ*_*j*_ and *γ*_*k*_, for SNPs *j* and *k*

where each SNP *j* is proposed according to a pre-calculated weight *w*_*j*_. The novelty of the proposal distribution is that we construct the weights *w*_*j*_’s based on the conditional analysis results. More specifically, we start by computing Bayes factors for each *cis*-SNP in a single SNP analysis, and compute a quantity

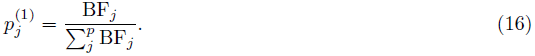

(Note that 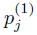 is proportional to the PIP for SNP *j* assuming only one eQTL in the *cis* region and a uniform prior inclusion probability). We then find the SNP with the maximum 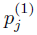 value, say SNP *k*. In the next round, we control for the genotype of SNP *k* and repeat the single SNP analysis to obtain 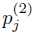, which mimics the conditional analysis of secondary *cis*-eQTL signals. Note that SNP *k* and the SNPs in LD will have single SNP Bayes factor close to 1 in this round. We again add the SNP with the maximum 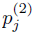 value into the control set. We repeat this procedure, with one additional SNP added into the control set in each round, until the maximum single SNP Bayes factor falls below a pre-defined threshold (we use 10 in practice). Suppose that the procedure ends in *t* iterations, we then compute the weight for each SNP using

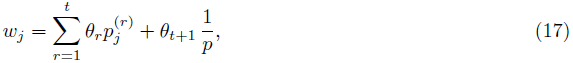

where the sequence *θ*_1_,…, *θ*_*t*+1_ forms a decreasing geometric series summing up to 1. The trailing 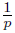 term in the weight calculation represents a uniform distribution on candidate *cis*-SNPs.

This particular proposal distribution is an extension of what is used in [37], and should be credited to Matthew Stephens (personal communication). Its theoretical backend is related to *sure-independence screening* proposed by [38] in variable selection context.

### S.2 Maximum Likelihood Inference of Enrichment Parameters

This section gives the technical details of MCMC-within-EM algorithm. Given the hierarchical model described in the main text, we are interested in performing maximum likelihood inference of enrichment parameter ***α***. Treating 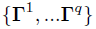 across all *q* genes as missing data, the complete data likelihood can be written as

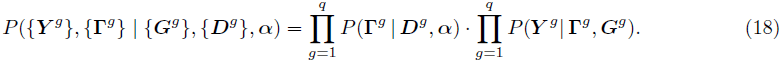

We apply an EM algorithm to find the MLE of ***α***. Because vector 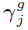 only takes values in {**0**, **1**}, using a loose notation, we represent vectors **0** and **1** with the corresponding binary scalar values. It then follows that

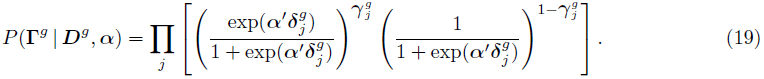

The complete data log-likelihood is given by

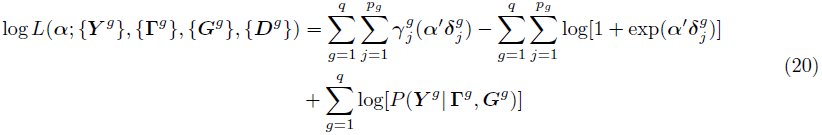

The EM algorithm initiates by an arbitrary value of ***α***, namely, ***α***^(1)^. In the E-step of *t*-th iteration, we compute

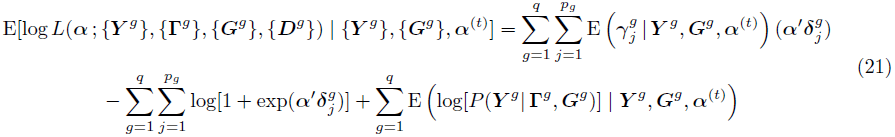

Note that the last term does not contain parameter ***α***. In the M-step of the *t*-th iteration, we find

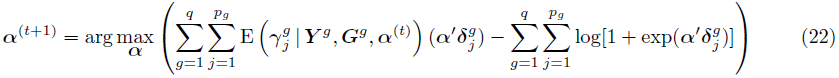

The objective function in (22) coincides with the log-likelihood function of a logsitic regression model treating each gene-SNP pair as an independent observation, however with the usual binary response variable replaced by the conditional expectations. By this connection, the maximization step can be carried out by fitting the corresponding modified logistic regression model treating conditional expectations as responses (i.e., via an iterative re-weighted least square algorithm). This also implies that in the E-step, it is only required to compute 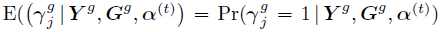, i.e., the PIP for each gene-SNP pair, which we obtain from the MCMC sampling.

To summarize, we outline the procedure of the MCMC-within-EM algorithm based on the above derivation as follows

1. At *t* = 1, initiate ***α*** = ***α***^(1)^
2. Compute prior Pr(**Γ**^*g*^ | ***D***^*g*^, ***α***^(*t*)^), and run MCMC algorithm for multiple *cis*-eQTL analysis for each gene *g*
3. Compute 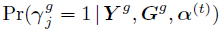 for each gene-SNP pair from the posterior samples
4. Find ***α***^(*t*+1)^ by fitting a logistic regression model treating 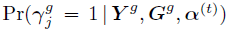 as response variable and {***D***^*g*^} as observed covariates
5. Repeat 2 to 4 until convergence

### S.3 Single Base-pair Resolution Annotation of Genetic Variants Predicted to Affect Transcription Factor Binding

The approach and validation of the annotation are detailed in [30] and here we provide a summary for the annotation method. The method used to develop the annotation is based on the CENTIPEDE approach that can predict TF activity from integrating sequence motifs together with functional genomics data. This approach gains the most information from high-resolution data such as DNase-seq or ATAC-seq [39]. In the original CENTIPEDE approach, the sequence models are pre-determined; e.g, k-mers or previously defined position weight matrix (PWM) models from databases such as TRANSFAC and JASPAR. However, many motif models were created with very few sample sequences obtained from known TF binding sites and do not represent the full spectrum of sequence variation that can be tolerated without affecting binding. To better capture this range, it is necessary to include motif instances that may not be a perfect match to the original PWM, but have evidence of binding in the human genome. In [30], we introduced a novel approach extending CENTIPEDE to re-adjust the sequence model for TF binding using only DNase-seq data in two steps:

#### Step 1: Initial CENTIPEDE scan and motif recalibration

After scanning the genome for motif matches (using 1949 seed motifs), we extracted DNase-seq data at these sites using 653 samples publicly available from the ENCODE and Roadmap Epigenomics projects. For each motif and only for this initial step, we used a reduced subset of motif matches that include the top 5,000 instances on the human genome; and up to 10,000 additional sequences in the human genome that do not have a high score. The low scoring motif instances were chosen from human sequences that have orthologous high scoring motif instances in the chimp or rhesus genome. We then applied the CENTIPEDE model to survey TF activity for each 1,272,697 tissue-TF pair. For each pair we then determined that the TF is active if the motifs instances that exhibit a CENTIPEDE footprint can be predicted from the PWM score (*z*-score > 5). Using this criterion, we determined that 1,891 TF motifs are active in at least one tissue. We then recalibrated the PWM model for each active motif using the sequences of all motif matches that have a DNase-seq footprint (CENTIPEDE posterior >0.99). Using this procedure, the probabilities of certain bases are readjusted, but the core part of the motif and its consensus sequence is largely maintained.

#### Step 2: Full genome CENTIPEDE scan and genetic variant analysis

Using these newly updated sequence models we scanned the human genome for all possible matches both to the reference and to alternate alleles from genetic variants cataloged in the 1000 Genomes (1KG) Project [27] and used the CENTIPEDE algorithm to assess the probability that each motif instance is bound by a TF. In this second step, we included all high and low scoring PWM matches down to a CENTIPEDE prior probability of binding of 10%. In this paper we focus only on SNPs found in CENTIPEDE footprints discovered in LCLs with a posterior probability > 0.99. In total about 600,000 SNPs are in LCL footprints, of which about half are predicted to strongly affect binding, affecting the prior odds of binding ≥ 20-fold (binding variants) based on the logistic sequence model hyper prior in the CENTIPEDE model.

### S.4 Automatic Clustering of Independent eQTL Signals from MCMC Output

We designed a hierarchical clustering based algorithm to automatically parse the MCMC output and recognize potentially multiple independent association signal clusters.

Let ***M*** denote the set of posterior models sampled by the MCMC algorithm. For each model *m*_*k*_ ∈ ***M***, we denote its posterior model probability by *p*_*m*_*k*__. We define a “distance” between SNP *i* and SNP *j* by

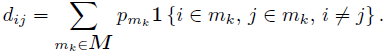

The above definition is the key to the algorithm. Every SNP has distance 0 to itself. For SNPs with high LD, they are inter-changeable of each other but almost never co-exist in a single posterior model, and consequently those SNPs have distances ≈ 0 with each other. On the other hand, SNPs representing independent signals do often co-exist in posterior models and have non-zero distance between each other. Consider a simple example with 3 SNPs: SNP 1 alone represents an independent signal and appears in all posterior models, SNP 2 and 3 are in high LD and jointly represent another independent signal. Suppose that from the posterior sampling, we observe posterior model [1, 2] and [1, 3] 40% and 60% of the time, respectively. The resulting pair-wise “distance” matrix based on our definition is then given by

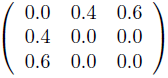

We then perform a hierarchical clustering based on the resulting pair-wise “distance” matrix constructed from the MCMC samples. By default, we choose the cluster number to be the maximum model size observed in all the posterior models. Based on the clustering result, we compute a cluster-level PIP by summing over the SNP-level PIPs within each inferred cluster. In the above toy example, by selecting cluster number *K* = 2, SNP 1 forms a cluster with the cluster-level PIP = 1.0 and SNP 2 and 3 form another cluster with cluster-level PIP = 1.0 as well.

It should be noted that our pair-wise distance measure is very similar to the commonly used Kullback-Liebler distance in measuring the independence between a pair of SNPs. However, our measure is more convenient to compute from the posterior model probabilities in the MCMC output. Neither our measure or the Kullback-Liebler distance is technically a well-defined distance metric. However in practice, the clustering algorithm works well with these pseudo distance measures.

We find that this algorithm performs reasonably well in practice. One of the most obvious advantage is that it automatically groups SNPs in high LD and recognizes clusters with high PIPs at cluster level. Nevertheless, we still view this algorithm as a heuristic tool to simply aid the post-hoc analysis. In addition, we find that checking the LD patterns from the genotype data for each inferred cluster can serve as a useful independent validation.

### S.5 Additional Details of Simulation Studies

In this section, we provide additional details on generating and analyzing simulated expression-genotype data in our simulation studies.

In the main text, we have detailed the procedure to assemble the genotype data from the GUEVADIS project. For each of the 1,500 simulated genes, we randomly sampled 1, 2, 3 or 4 regions to harbor a causal eQTL with probabilities 0.40, 0.30, 0.20 and 0.10, respectively. (For example, with probability 0.20, the simulated gene contain 3 independent eQTL signal.) Once the number of independent signals was determined, we randomly selected a single causal SNP as the causal SNP from each eQTL region according to a discrete uniform distribution.

For each causal SNP, we simulated its effects in the five populations according to the following scheme. We first generated a mean effect from 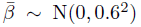, then the eQTL effect of the causal SNP for each population was subsequently drawn from the distribution 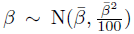. With this procedure, the resulting eQTL effects are highly correlated across populations however with some low levels of heterogeneity. It is worth pointing out that this generating model is different from the model that we used for the Bayesian analysis. Finally, we generated the expression phenotype separately in each population using the linear model (2.1), with the additional random error vector simulated from N(0,***I***).

To perform single SNP analysis on the simulated data set, we carried out a fixed effect meta-analysis procedure. More specifically, for SNP *j* in population group *i*, we computed a *z*-score 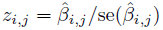 by fitting a simple linear regression model. We then computed a fixed effect meta-analysis test statistic

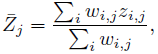

where the weight is obtained by 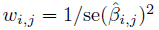. The variance of 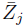 is calculated by

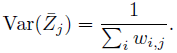

For each SNP, we computed an overall *z*-value and obtained its corresponding *p*-value.

The core procedure for the conditional analysis in each round is similar to the above descried single SNP analysis. However, instead of fitting a simple linear regression model, we fit a multiple regression model controlling for the top associated SNPs identified from the previous rounds. We started the procedure with an empty set of SNPs to be controlled for and halted the procedure until the most significant fixed effect meta-analysis *p*-value is larger than the pre-defined threshold.

## Supporting Information

### Supplementary Data

**S1 Dataset**. List of gene-SNP pairs displaying highly heterogeneous effects across populations flagged by single SNP analysis.

**S2 Dataset**. Complete enrichment estimates from the enrichment analysis of eQTL signals with respect to distance to TSS (200 bins) and TF binding site annotations.

## References

1. Nica AC, Montgomery SB, Dimas AS, Stranger BE, Beazley C, et al. (2010) Candidate causal regulatory effects by integration of expression QTLs with complex trait genetic associations. PLoS genetics 6: e1000895.

2. Nicolae DL, Gamazon E, Zhang W, Duan S, Dolan ME, et al. (2010) Trait-associated SNPs are more likely to be eQTLs: annotation to enhance discovery from GWAS. PLoS Genetics 6: e1000888.

3. Hao K, Bossé Y, Nickle DC, Paré PD, Postma DS, et al. (2012) Lung eQTLs to help reveal the molecular underpinnings of asthma. PLoS genetics 8: e1003029.

4. GTEx Consortium (2013) The genotype-tissue expression (gtex) project. Nature Genetics 45: 580–585.

5. Dimas AS, Deutsch S, Stranger BE, Montgomery SB, Borel C, et al. (2009) Common regulatory variation impacts gene expression in a cell type–dependent manner. Science 325: 1246–1250.

6. Maranville JC, Luca F, Richards AL, Wen X, Witonsky DB, et al. (2011) Interactions between glucocorticoid treatment and cis-regulatory polymorphisms contribute to cellular response phenotypes. PLoS genetics 7: e1002162.

7. Barreiro LB, Tailleux L, Pai AA, Gicquel B, Marioni JC, et al. (2012) Deciphering the genetic architecture of variation in the immune response to mycobacterium tuberculosis infection. Proceedings of the National Academy of Sciences 109: 1204–1209.

8. Raj T, Rothamel K, Mostafavi S, Ye C, Lee MN, et al. (2014) Polarization of the effects of autoimmune and neurodegenerative risk alleles in leukocytes. Science 344: 519–523.

9. Brown CD, Mangravite LM, Engelhardt BE (2013) Integrative modeling of eQTLs and cisregulatory elements suggests mechanisms underlying cell type specificity of eQTLs. PLoS Genetics 9: e1003649.

10. Birney E, Stamatoyannopoulos JA, Dutta A, Guigó R, Gingeras TR, et al. (2007) Identification and analysis of functional elements in 1% of the human genome by the encode pilot project. Nature 447: 799–816.

11. Pique-Regi R, Degner JF, Pai AA, Gaffney DJ, Gilad Y, et al. (2011) Accurate inference of transcription factor binding from dna sequence and chromatin accessibility data. Genome research 21: 447–455.

12. Hoffman MM, Ernst J, Wilder SP, Kundaje A, Harris RS, et al. (2012) Integrative annotation of chromatin elements from encode data. Nucleic acids research: gks1284.

13. Liang L, Morar N, Dixon AL, Lathrop GM, Abecasis GR, et al. (2013) A cross-platform analysis of 14,177 expression quantitative trait loci derived from lymphoblastoid cell lines. Genome research 23: 716–726.

14. Flutre T, Wen X, Pritchard J, Stephens M (2013) A statistical framework for joint eQTL analysis in multiple tissues. PLoS Genetics 9: e1003486.

15. Wen X (2014) Bayesian model selection in complex linear systems, as illustrated in genetic association studies. Biometrics 70: 73–83.

16. Sul JH, Han B, Ye C, Choi T, Eskin E (2013) Effectively identifying eQTLs from multiple tissues by combining mixed model and meta-analytic approaches. PLoS genetics 9: e1003491.

17. Gaffney DJ, Veyrieras JB, Degner JF, Pique-Regi R, Pai AA, et al. (2012) Dissecting the regulatory architecture of gene expression QTLs. Genome Biol 13: R7.

18. Veyrieras JB, Kudaravalli S, Kim SY, Dermitzakis ET, Gilad Y, et al. (2008) High-resolution mapping of expression-QTLs yields insight into human gene regulation. PLoS genetics 4: e1000214.

19. Lee SI, Dudley AM, Drubin D, Silver PA, Krogan NJ, et al. (2009) Learning a prior on regulatory potential from eQTL data. PLoS genetics 5: e1000358.

20. Lappalainen T, Sammeth M, Friedländer MR, T Hoen PA, Monlong J, et al. (2013) Transcriptome and genome sequencing uncovers functional variation in humans. Nature 501: 506–511.

21. Wen X, Stephens M (2014) Bayesian methods for genetic association analysis with heterogeneous subgroups: From meta-analyses to gene–environment interactions. The Annals of Applied Statistics 8: 176–203.

22. Morris AP (2011) Transethnic meta-analysis of genomewide association studies. Genetic epidemiology 35: 809–822.

23. Marigorta UM, Navarro A (2013) High trans-ethnic replicability of gwas results implies common causal variants. PLoS genetics 9: e1003566.

24. Li G, Shabalin AA, Rusyn I, Wright FA, Nobel AB (2013) An empirical bayes approach for multiple tissue eQTL analysis. arXiv preprint arXiv:13112948.

25. Yang J, Ferreira T, Morris AP, Medland SE, Madden PA, et al. (2012) Conditional and joint multiple-snp analysis of gwas summary statistics identifies additional variants influencing complex traits. Nature Genetics 44: 369–375.

26. Coviello AD, Haring R, Wellons M, Vaidya D, Lehtimäki T, et al. (2012) A genome-wide association meta-analysis of circulating sex hormone–binding globulin reveals multiple loci implicated in sex steroid hormone regulation. PLoS genetics 8: e1002805.

27. 1000 Genomes Project Consortium, et al. (2012) An integrated map of genetic variation from 1,092 human genomes. Nature 491: 56–65.

28. Gilad Y, Rifkin SA, Pritchard JK (2008) Revealing the architecture of gene regulation: the promise of eQTL studies. Trends in Genetics 24: 408–415.

29. Degner JF, Pai AA, Pique-Regi R, Veyrieras JB, Gaffney DJ, et al. (2012) DNase I sensitivity QTLs are a major determinant of human expression variation. Nature 482: 390–394.

30. Moyerbrailean GA, Harvey CT, Kalita CA, Wen X, Luca F, et al. (2014) Are all genetic variants in dnase i sensitivity regions functional? bioRxiv: 007559.

31. Harrow J, Frankish A, Gonzalez JM, Tapanari E, Diekhans M, et al. (2012) Gencode: the reference human genome annotation for the encode project. Genome research 22: 1760–1774.

32. Pickrell JK, Marioni JC, Pai AA, Degner JF, Engelhardt BE, et al. (2010) Understanding mechanisms underlying human gene expression variation with rna sequencing. Nature 464: 768–772.

33. Stegle O, Parts L, Piipari M, Winn J, Durbin R (2012) Using probabilistic estimation of expression residuals (PEER) to obtain increased power and interpretability of gene expression analyses. Nature protocols 7: 500–507.

34. Wen X (2013) Robust bayesian FDR control with bayes factors. arXiv preprint arXiv:13113981.

35. Newton MA, Noueiry A, Sarkar D, Ahlquist P (2004) Detecting differential gene expression with a semiparametric hierarchical mixture method. Biostatistics 5: 155–76.

36. Fairfax BP, Humburg P, Makino S, Naranbhai V, Wong D, et al. (2014) Innate immune activity conditions the effect of regulatory variants upon monocyte gene expression. Science 343: 1246949.

37. Guan Y, Stephens M, et al. (2011) Bayesian variable selection regression for genome-wide association studies and other large-scale problems. The Annals of Applied Statistics 5: 1780–1815.

38. Fan J, Lv J (2008) Sure independence screening for ultrahigh dimensional feature space. Journal of the Royal Statistical Society: Series B (Statistical Methodology) 70: 849–911.

39. Buenrostro JD, Giresi PG, Zaba LC, Chang HY, Greenleaf WJ (2013) Transposition of native chromatin for fast and sensitive epigenomic profiling of open chromatin, dna-binding proteins and nucleosome position. Nature methods.

